# FluoEM: Virtual labeling of axons in 3-dimensional electron microscopy data for long-range connectomics

**DOI:** 10.1101/340802

**Authors:** Florian Drawitsch, Ali Karimi, Kevin M. Boergens, Moritz Helmstaedter

## Abstract

Volume electron microscopy (3D EM) has enabled the dense reconstruction of neuronal circuits in datasets that are so far about a few hundred micrometers in extent. In mammalian brains, most neuronal circuits are however highly non-local, such that a large fraction of the synapses in such a volume of neuropil originates from distant projection sources. The labeling and identification of such long-range axonal inputs from multiple sources within a densely reconstructed EM dataset has been notoriously difficult because of the limited color label space of EM. Here, we present FluoEM, a set of experimental and computational methods that allows the identification of multi-color fluorescently labeled axons in dense EM data without the need for artificially introduced fiducial marks or direct label conversion for EM. The approach is based on correlated imaging of the tissue and computational matching of neurite reconstructions, amounting to a virtual color labeling of axons in dense EM circuit data. We show that the identification of fluorescent light-microscopically (LM) imaged axons in 3D EM data from mouse cortex is faithfully possible as soon as the EM dataset is about 40-50 μm in extent, relying on the unique trajectories of axons in dense mammalian neuropil. The method is exemplified for the identification of longdistance axonal input into layer 1 of the mouse cerebral cortex.

## Introduction

The dense reconstruction of neuronal circuits has become an increasingly realistic goal in the neurosciences thanks to developments in large-scale 3D EM (Denk and Horstmann 2004, Bock, Lee et al. 2011, Hayworth, Xu et al. 2015, Kasthuri, Hayworth et al. 2015, Xu, Hayworth et al. 2017) and circuit reconstruction techniques (Saalfeld, Cardona et al. 2009, Boergens, Berning et al. 2017). First locally dense neuronal circuits have been mapped (Helmstaedter, Briggman et al. 2013, Kasthuri, Hayworth et al. 2015, Berck, Khandelwal et al. 2016, Wanner, Genoud et al. 2016). However, in mammalian nervous systems, any given local neuronal circuit receives numerous synaptic inputs from distant neuronal sources. In most layers of the mammalian cortex, for example, the fraction of distal input synapses in a dense local circuit is estimated to comprise 70-90% of all synapses (Stepanyants, Martinez et al. 2009). Similarly, subcortical structures as the striatum or amygdala (Fig. 1a) receive the vast majority of their inputs from distal projection sources. Uncovering the synaptic logic of such multi-source circuits together with the local dense neuronal connectivity requires the identification of multiple input sources in the same connectomic experiment. Since 3D EM is still limited to volumes that do not routinely encompass entire mammalian brains, the origin of a large fraction of input synapses remains thus unidentified in current high-resolution connectomic experiments in mammals.

**Figure 1.**
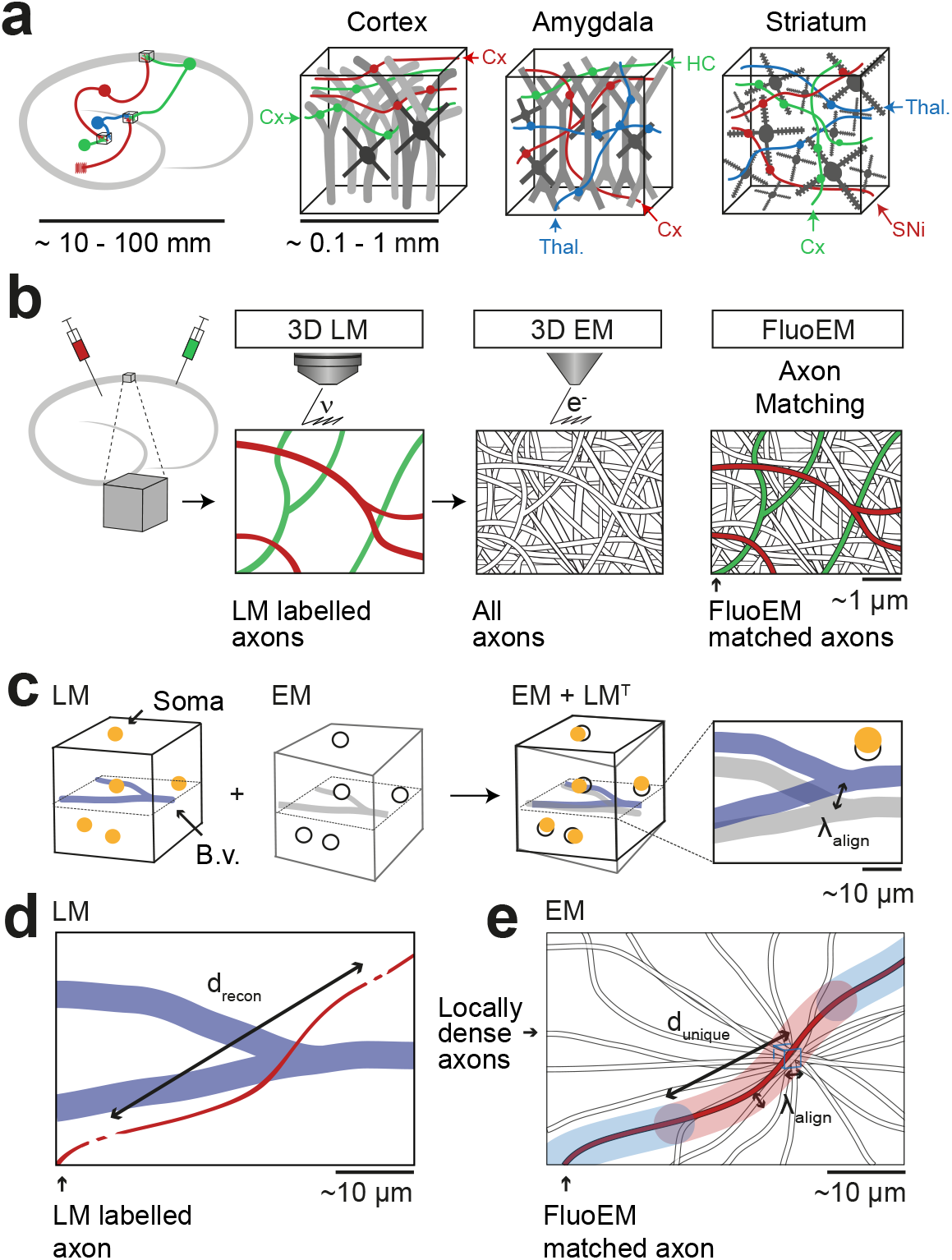
FluoEM applications and concept. (**a**) Most local circuits in mammalian brains can be mapped densely by modern 3D EM (gray neurons), but the source of a majority of the relevant input axons remain unidentified (colored), because the projections arise from distal brain regions (see sketch on the left). Examples show upper layers of cortex, Amygdala and Striatum with dominant input from other cortices (Cx), Thalamus (Thal.), Hippocampus (HC), Substantia Nigra (SNi). Deciphering the connectomic logic of these inputs within a densely imaged neuronal circuit requires parallel axonal labels in a single EM experiment. Mapping color-encoded information on multiple axonal origins from LM data into 3D EM data useable for connectomic reconstruction in mammalian nervous tissue is the goal of the presented method. (**b**) A brain tissue sample containing fluorescently labeled axons is volume imaged first using confocal laser scanning microscopy (LM) and then using 3D Electron Microscopy (EM). The fluorescently labeled axons (red and green) contained in the LM dataset are then matched to the corresponding axons in the EM dataset (black and white) solely based on structural constraints (see b-d), without chemical label conversion or artificial fiducial marks. (**c**) Coarse LM-EM volume registration based on intrinsic fiducials such as blood vessels (blue) and somata (yellow) can be routinely achieved at a registration precision *λ*_align_ of about 5-10 μm. (d) The path length *d_recon_* over which fluorescently labeled axons (green and red) in a volume LM dataset are clearly reconstructable depends mostly on the labeling density in the respective fluorescence channel. Blue, blood vessels (B.v.). (e) Axons (grey) traversing a common bounding box (blue) of extent *λ*_align_ become increasingly distinguishable with increasing distance from the common origin. After a distance *d_unique_*, a given axonal trajectory is unique, i.e. no other axon from the center bounding box is still closer than *λ_align_*. Since 3D EM allows the reconstruction of all axons traversing a given bounding box, *d_unique_* can be determined (see Fig. 2). If axons can be reconstructed at the LM level for a path length *d_recon_* (c) that is similar to or larger than the typical axonal uniqueness length *d_unique_* (d), the FluoEM approach can in principle be successful.

At the same time, the labelling of multiple neuronal populations and their axonal projections by fluorescent dyes visible in LM is routine (Livet, Weissman et al. 2007, Mao, Kusefoglu et al. 2011, D’Souza, Meier et al. 2016). Combining light-microscopic fluorescent labels in axons from multiple sources with large-scale 3D EM would be ideal. Such direct correlative light- and electron microscopy (CLEM) has been successful for high-resolution subcellular label localization (Agronskaia, Valentijn et al. 2008, Murphy, Narayan et al. 2011, Liv, Zonnevylle et al. 2013) and synaptic identification (Micheva and Smith 2007, Rah, Bas et al. 2013), but can so far not be combined with 3D EM at both resolution and scale allowing for dense large-scale circuit mapping in mammalian neuropil.

Similarly, en-bloc immunolabeling in conjunction with EM is limited to thin tissue slices (Faulk and Taylor 1971, Pallotto, Watkins et al. 2015). Approaches to interlace EM and LM imaging on alternating tissue slices, though successfully applied in invertebrates (Shahidi, Williams et al. 2015), impede dense EM-based circuit reconstruction in mammalian nervous tissue. Rather, methods to obtain electron-dense labels in a subset of neurons have been developed over the decades based on induced axonal degeneration (Gray and Hamlyn 1962, Colonnier 1964), introduction of HRP (Hamos, Van Horn et al. 1985, Horikawa and Armstrong 1988, Anderson, Douglas et al. 1994, Anderson, Douglas et al. 1994, Markram, Lübke et al. 1997, Costa and Martin 2011) or fluorophores (Maranto 1982, Grabenbauer, Geerts et al. 2005, Knott, Holtmaat et al. 2009) into a subset of neurons which could be utilized to create electron contrast via enzymatic or photo-oxidative DAB polymerization. More recent improvements enabled the genetically targeted expression of electron-dense labels in diverse cellular compartments, potentially allowing to encode multiple label classes in a single EM experiment (Shu, Lev-Ram et al. 2011, Martell, Deerinck et al. 2012, Atasoy, Betley et al. 2014, Lam, Martell et al. 2015, Joesch, Mankus et al. 2016). However, generating multiple electron-dense label classes in the very same experiment while ensuring ultrastructural preservation is challenging and to our knowledge has been accomplished in drosophila (Lin, Luo et al. 2016) but so far not in mammals. Furthermore, the number of cellular compartments usable for such axonal identification may be rather limited.

Methods to directly co-register light microscopic labels in axons and dendrites with 3D EM data (Bishop, Nikić et al. 2011, Maco, Holtmaat et al. 2013) have been restricted to local volumes of up to 10 μm on a side, using LM-burnt fiducial marks visible in EM; these do not scale to the large volumes required for neuronal circuit reconstruction (sized hundreds of micrometers per dimension, i.e. at least 1000-fold larger). Direct co-alignment between fluorescently labeled cell bodies and EM data has been routinely applied (Bock, Lee et al. 2011, Briggman, Helmstaedter et al. 2011, Lee, Bonin et al. 2016, Tsang, Bushong et al. 2018), but provides LM-to-EM alignment only at a coarse scale that does not allow for the identification of single axons.

Apart from the experimental limitations currently hampering the use of multi-color labels in large 3D EM data, current computational approaches are challenged when attempting submicron registration precision of neurites in volumes required for connectomics (∼10^6^ μm^3^ or larger). Previously proposed multi-modal graph-based neurite registration (Serradell, Glowacki et al. 2012, Fua and Knott 2015, Serradell, Pinheiro et al. 2015) can in principle be used for LM-to-EM morphological matching, but did not yet scale to graphs of thousands or tens of thousands of nodes as required for connectomics.

Thus, a method that could directly employ the full multi-color space of LM in large 3D tissue samples while enabling dense EM-based circuit reconstruction would be ideal. Here we report FluoEM, a set of experimental and computational tools to achieve this goal. Instead of directly attempting alignment of LM-labeled axons to the EM dataset at the nanometer scale, we exploit the availability of locally dense neurite reconstructions in current volume EM. Here, the alignment problem can be reformulated as identifying the most likely axon (out of all axons, reconstructed in EM) that best explains an LM-imaged axonal fluorescence signal. We show that this is a well-constrained problem based on the actual geometry of axonal fibers in nerve tissue from mouse cerebral cortex. We report the experimental methods allowing for subsequent correlative imaging of samples in LM and EM and the computational tools to register the labeled LM axons to their EM correspondences. We exemplify our method for long-range inputs to layer 1 of mouse cortex.

## RESULTS

The FluoEM workflow (Fig. 1b) comprised the following steps: acquisition of a 3D fluorescence-LM dataset from a given piece of tissue; 3D imaging of that very same piece of tissue in the electron microscope; reconstruction of the fluorescently labeled axons in the LM data; reconstruction of locally all axons in small subvolumes of the EM data; computational determination of the most likely EM axons that explain the LM signal.

In a first step, we used blood vessels as intrinsic fiducials in the neuropil to obtain a coarse co-registration between LM and EM (Fig. 1b). Coarse LM-EM registration based on cell bodies and blood vessels has been successfully performed before (Bock, Lee et al. 2011, Briggman, Helmstaedter et al. 2011) and is expected to yield an alignment precision *λ_align_* of about 5-10 micrometers (Fig. 1c). We next had to determine the typical length of faithful 3D axon reconstruction *d_recon_* in fluorescence data (Fig. 1d). Then, the problem of matching EM axons to LM data can be phrased as follows (Fig. 1e): Given all axons that traverse a cube of edge length *λ_align_* in the EM data, will the trajectory of these axons become unique on a length scale *d_unique_* that is smaller than or comparable to the LM-reconstruction length *d_recon_: d_unique_(*λ_align_*) ≤ d_recon_*? In other words: What is the travel length *d_unique_* after which all axons from a center cube of size *λ_align_* will have no other of these axons closer than *λ_align_*?

In order to understand whether this approach could work, we had to measure *d_recon_* in LM data with realistic labeling density and determine *d_unique_* for a realistic *λ_align_* in EM data. These measurements are reported next, followed by a proof-of-principle coregistration experiment and results from the virtual labeling of axons in EM data.

### Reconstruction length and axon uniqueness

We first performed a fluorescence imaging experiment (Fig. 2a-c) in which we injected adeno-associated viruses expressing fluorescent markers into the primary motor cortex (M1; virus expressing the green marker eGFP) and the secondary somatosensory cortex (S2; virus expressing the red marker tdTomato) of a 28-day old mouse, followed by transcardial perfusion and tissue extraction at 46 days of age. Using a confocal laser scanning microscope we then acquired a dataset sized (828 × 828 × 75) μm^3^ at a voxel size of (104 × 104 × 444) nm^3^ from layer 1 of a part of cortex in which both of these projections converged (located to lateral parietal association cortex (LPtA), Fig. 2b). We then 3D-reconstructed all axons traversing a (40 × 40 × 48) μm^3^ bounding box in this LM image dataset using our annotation tool webKnossos (Boergens, Berning et al. 2017) and measured the length over which the axons were faithfully reconstructable as judged by expert annotators (for this we asked the annotators to stop reconstructing whenever they encountered a location of any ambiguity, the loss of axon continuity or an unclear branching). The reconstruction length *d_recon_* per axon was 70 ± 44 μm (mean ± s.d., n=46 axons) for the green and 134 ± 62 μm (n=42 axons) for the red fluorescence channel. 95% of axons were reconstructable above 20 μm length, some for up to 200 μm (Fig. 2c, independent reconstruction by two experts).

**Figure 2.**
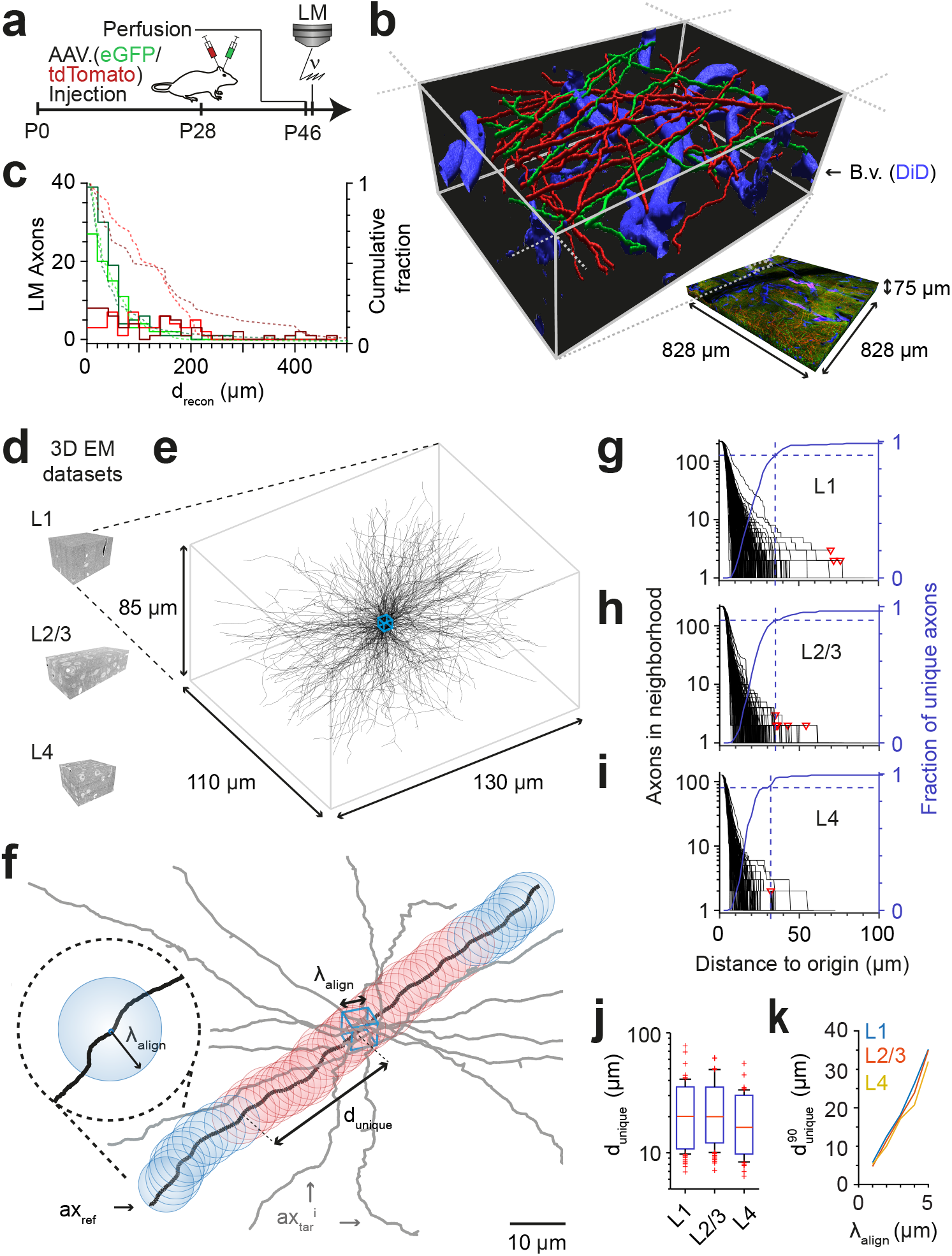
FluoEM proof-of-principle measurements. (**a-c**), Measurement of the length *d_recon_* over which axons in LM fluorescence data are faithfully reconstructable (see Fig. 1d). (**a**) Experimental timeline: Injection of a fluorescent-protein (FP) expressing adeno-associated virus at postnatal day 28 (AAV.eGFP into M1 cortex, AAV.tdTomato into S2 cortex of a wild-type C57BL/6j mouse); transcardial perfusion, fixation, sample extraction from cortical L1 at postnatal day 46 followed by confocal volume imaging. (**b**) Rendering of a subvolume of the 3-channel confocal image stack showing axonal long-range projections from M1 cortex (green), S2 cortex (red) as well as stained blood vessels (blue). (**c**) Histogram of the reconstructable pathlength (d_recon_) for the fluorescently stained long-range axons (red, green) in the example LM dataset (b) based on two independent expert annotations (light and dark colors; median *d_recon_* of 28 and 33 μm (green channel, n=115 and 88) and 88 and 107 μm (red channel, n=56 and 47) for the two annotators, respectively). Dashed lines: cumulative axon fraction per annotator and channel. Note that annotators were explicitly asked to stop reconstructions whenever any ambiguity or uncertainty occurred, thus biasing these reconstructions to shorter length. (**d-k**), EM-based measurements of the length *d_unique_* over which the trajectory of axons becomes unique in several cortical layers. (**d**) Renderings of three 3D EM datasets in layers 1, 2/3 and 4 of mouse cortex. (**e**) Skeleton reconstructions in the L1 3D EM dataset of all 220 axons (black) that traversed a center bounding box (blue) of edge length *λ_align_ = 5 μm*. (**f**) Determination of the uniqueness length *d_unique_* for a given axon (black) by counting the number of other axons (gray) that were within the same seeding volume (blue box) and remain within a distance of no more than *λ_align_* from the reference axon for increasing distances *d* from the center box. *d_unique_* was defined as the Euclidean distance from the center box at which no other axon persistently was at less than *λ_align_* distance from the reference axon (red: at least one neighboring axon, blue: no more neighboring axon). (**g**) Axonal uniqueness length *dunique* in cortical L1 for all n=220 axons from the center bounding box (see e). Number of neighboring axons for each axon (black) and the combined fraction of unique axons (blue) over Euclidean distance from the bounding box center. Note that at 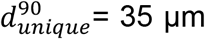 the trajectory of 90% of all axons has become unique (dashed blue line). For three of 220 axons, one or two other axons remained in *λ_align_* proximity within the dataset (red triangles). (**h,i**) Axonal uniqueness length in two additional 3D EM datasets from L2/3 (h, n=207 axons) and L4 (i, n=128 axons), labels as as in g. (**j**) Box plot of uniqueness length per axon for cortical layers 1-4 (from g-i). Boxes indicate 10,90-percentiles, whiskers indicate 5,95-percentiles, red crosses indicate outliers. (**k**) Effect of the coarse alignment precision *λ*_align_ on axonal uniqueness length 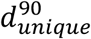 (resampled for smaller *λ_align_*=1,2,3,4 μm, see Methods).

We then measured the length *d_unique_* over which the trajectory of axons in cortex becomes unique, assuming a macroscopic alignment precision of *λ_align_* = 5 μm (Fig. 2d-k). For this, we acquired a 3D EM dataset sized (130 × 110 × 85) μm^3^ at a voxel size of (11.24 × 11.24 × 30) nm^3^ from layer 1 of LPtA cortex (Fig. 2d,e) using SBEM (Denk and Horstmann 2004) and reconstructed all n = 220 axons that traversed a cubic bounding box of size (5 × 5 × 5) μm^3^ in the center of the dataset (Fig. 2e, blue box). We then analyzed for each axon how many of the other 219 axons were still within *λ_align_* proximity at a Euclidean distance *d* from the center cube (Fig. 2f). For a given axon, the uniqueness length was then computed as the distance after which no other axon was in the *λ_align_*-surround. Fig. 2g shows the number of remaining axons after distance *d* for a dense reconstruction of axons in L1 of mouse cortex. The uniqueness length per axon was 22 ± 10 μm (mean ± s.d., range 7 – 78 μm, n=220 axons); after 31, 35, 41 and 69 μm, 85%, 90%, 95% and 98% of axons had no other axon in their proximity, respectively. We repeated this analysis in two additional 3D EM datasets obtained from layers 2/3 and 4 of mouse somatosensory cortex (datasets 2012-09-28_ex145_07x2 and 2012-11-23_ex144_st08x2, KM Boergens & MH, unpublished, Figs 2h,i). Here, uniqueness lengths per axon were 21 ± 9 μm (mean ± s.d., range 7 – 62 μm, n=214, L2/3) and 17 ± 8 μm (mean ± s.d., range 6-55 μm, n=128, L4), with 90% of axons unique after 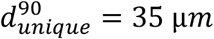 in L2/3 and 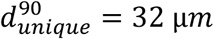 in L4 (Fig. 2j, 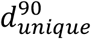 was indistinguishable between layers 1, 2/3 and 4, Fig. S1).

Importantly, when comparing 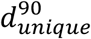 in layer 1 as determined in 3D EM data to the reconstruction lengths *d_recon_* obtained in the fluorescence datasets (Fig. 2c), we find that about 45% (green channel) and about 80% (red channel) of the faithfully LM-reconstructed axonal stretches were at least of length 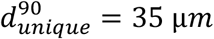. Since for a first LM-EM co-registration a set of the longest LM-reconstructable axons can be used, these data strongly suggested that the FluoEM approach for matching-based co-alignment between LM and EM reconstructed axons could in principle be successful. Furthermore, we expected the uniqueness length 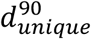 to depend on the macroscopic alignment precision *λ_align_* achieved by the blood-vessel based coarse pre-registration of the LM and EM datasets (Fig. 2k). When re-analyzing our dense axonal reconstructions in the 3D EM dataset from L1 for smaller *λ_align_*, we find that the uniqueness length strongly decreased for smaller *λ_align_* (to 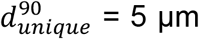 at *λ_align_* = 1 μm; 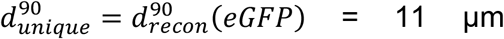 at *λ_align_* = 2 μm; 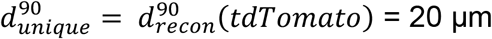 at *λ_align_* = 3 μm).

### Correlated LM and 3D EM dataset acquisition

We then set out to perform correlated 3D LM-EM imaging for the application of FluoEM (Fig. 3). For this, we continued the experiment described above (Fig. 2a) and acquired an additional fluorescence channel to image blood vessels (during perfusion, the rinsing solution was supplemented with DiD, a lipophilic dye for blood vessel staining with fluorescence in the far red spectrum (around 650 nm)). Then we applied our standard volume EM staining protocol to the tissue block (Hua, Laserstein et al. 2015) and imaged the sample using SBEM (Denk and Horstmann 2004) (Fig. 3a). We first acquired EM images with a large field of view (dataset dimensions (1500 × 1000 × 10) μm^3^) at lower resolution of (270 × 270 × 200) nm^3^ to identify the location of the LM-imaged dataset relative to the EM imaged sample (Fig. 3b,c). For this, we used the pattern of superficial blood vessels as coarse alignment fiducials (Fig. 3d,e). We then acquired a high-resolution 3D EM dataset sized (130 × 110 × 85) μm^3^ at a voxel size of (11.24 × 11.24 × 30) nm^3^ in a region previously imaged using LM with suitable fluorescent labeling density (Fig. 3c,g). In both the high-resolution EM dataset and the corresponding LM volume we then reconstructed the blood vessels (Fig. 3h,i) and annotated their branch points as control points for fitting a coarse affine transformation AT_BV_ from the LM to the high resolution EM dataset (Figs. 3h-n, n=7 control points). To estimate the precision of this coarse alignment, we measured its residuals (Fig. 3o; 2.7 ± 1.1 μm, mean ± s.d., n=7, median = 2.7 μm). Since this coarse alignment precision was well in the range previously assumed for the analysis of axonal uniqueness (*λ_align_* ≈ 5 μm, Fig. 3g-k), we then attempted a first LM-to-EM matching.

**Figure 3.**
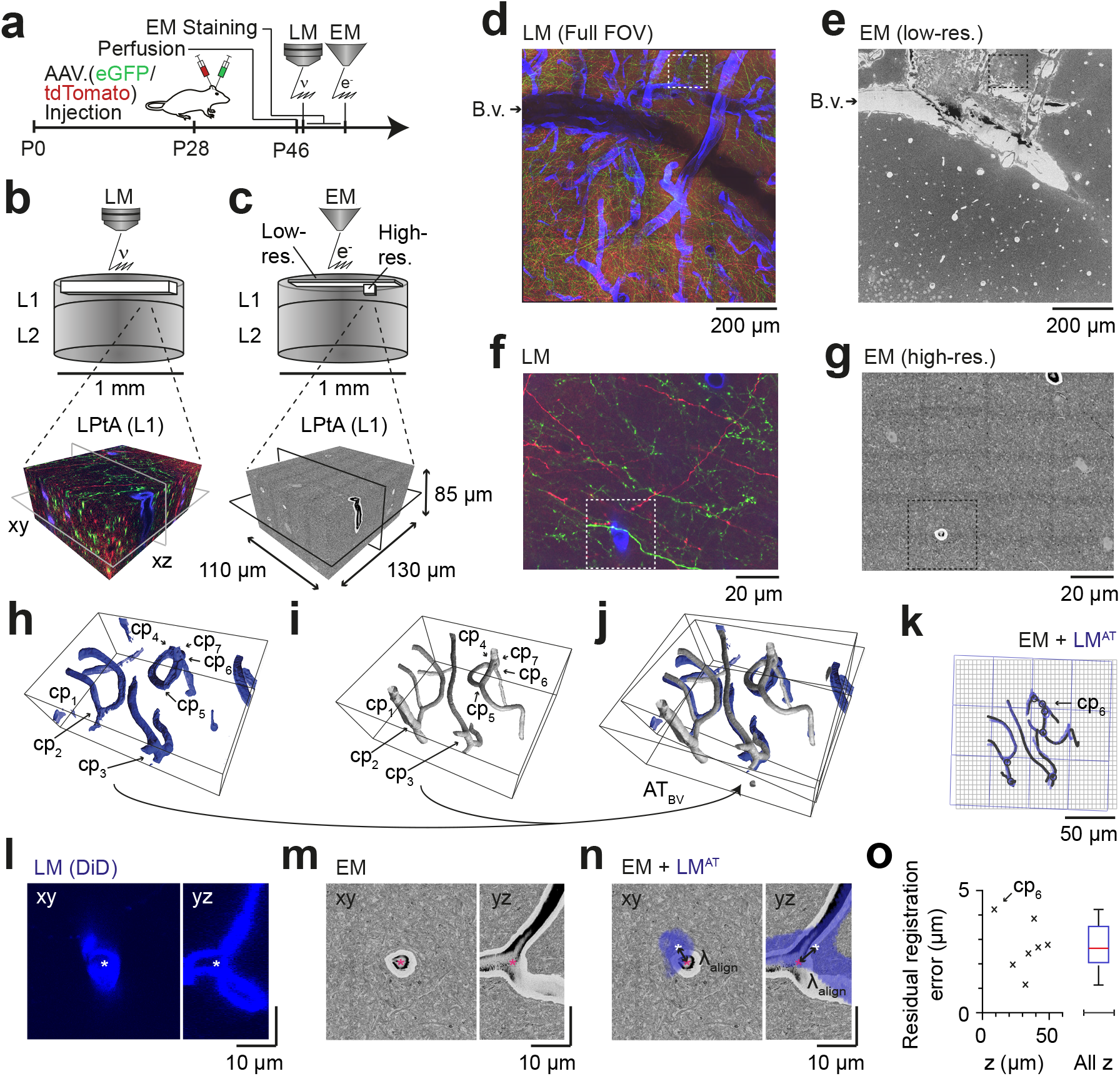
Coarse LM-EM registration. (**a**) Experimental timeline for FluoEM experiment (continued from Fig. 2a). After LM imaging was concluded the sample was immediately stained for EM and embedded, followed by 3D EM imaging. (**b,c**) Sketch of sequentially acquired 3D LM and EM datasets from cortical layer 1 (L1). Fluorescence channels: green (axonal projections from M1 cortex, eGFP); red (axonal projections from S2 cortex, tdTomato); blue (blood vessels, DiD). Volume and voxel size of EM datasets: (1500 × 1000 × 10) μm^3^ and (700 × 700 × 200) nm^3^ (Low resolution EM dataset), (130 × 110 × 85) μm^3^ and (11.24 × 11.24 × 30) nm^3^ (High resolution EM dataset). (**d,e**) Overview images of the LM (d) and low resolution EM (e) datasets. Dashed rectangle indicates the position of the high-resolution EM dataset (g, see c for relative positions of datasets). B.v., blood vessel. Note the corresponding pattern of surface vessels in d,e. (**f,g**) High-resolution images at corresponding locations in LM (f) and EM (g). Dashed rectangles indicate regions shown in l-n. (**h,i**), Rendering of blood vessel segmentations in LM (h) and EM (i) and their characteristic bifurcations used as control point pairs (cp_1-7_) to constrain a coarse affine transformation (AT_BV_). (**j**) Overlay of the EM and LM^AT^ blood vessel segmentations registered using AT_BV_. (**k**) Registered blood vessels (lines) and control points (circles) overlaid with EM (black) and transformed LM (blue) coordinate grids. (l-n) LM (l) and EM (m) image planes (xy) and reslice (yz) showing an exemplary blood vessel bifurcation (asterisk indicates cp6, see h-k) and the overlay (n) of the affine transformed LM blood vessel segmentation with EM. Arrows indicate registration residual, a measure of the coarse alignment precision *λ*_align_. (**o**) Alignment precision *λ*_align_ reported as residuals of control point alignment along dataset depth (z) (mean: 2.7 ± 1.1 μm, median: 2.7 μm) after affine transformation *AT_BV_*.

### LM-to-EM axon matching

We chose an axon next to a blood vessel in the LM data (Fig. 4a). The axon was skeleton-reconstructed at the LM level (Fig. 4b) using webKnossos (Boergens et al., 2017). We then transformed this LM reconstruction (path length of 105 μm) to the EM dataset using the coarse transformation AT_BV_ based on blood vessel correspondence as determined before (Fig. 4c). Evidently, based solely on this coarse transformation, an identification of the corresponding axon at the EM level was impossible (Fig. 4d). Rather, we then defined a cubic region sized (5 μm)^3^ in the EM data next to the blood vessel at which the LM reconstruction had been seeded (Fig. 4e,f) and reconstructed all axons that traversed that region in the EM dataset (Fig. 4g, n=193 axons). We then used a spherical search kernel with 5 μm radius (Fig. 2f) to determine which of the 193 EM-reconstructed axons were within that surround of the transformed LM-reconstructed axon along all of its trajectory (Fig. 4h,i). Beyond a distance of 42 μm along the transformed LM axon, only one candidate EM-reconstructed axon remained persistently within the search kernel (Fig. 4i). So, in fact, for this example fluorescent axon, we had found a corresponding axon at the EM level that matched its trajectory.

**Figure 4.**
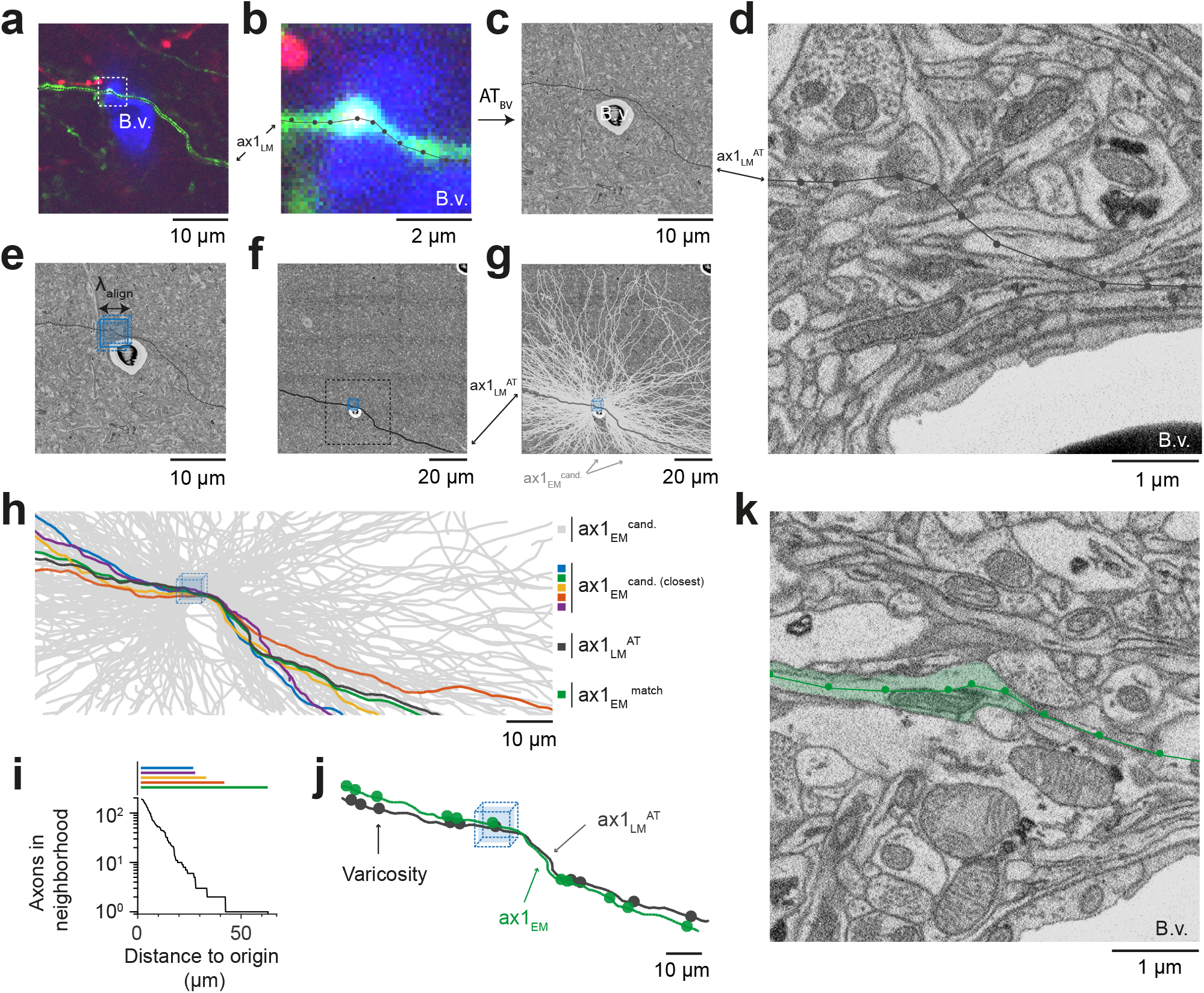
Initial LM-to-EM axon matching. (**a,b**) LM image plane showing an axon (green) overlaid with its LM-based skeleton reconstruction (ax1_LM_, black) and a nearby blood vessel bifurcation (B.v., blue). Dashed rectangle, region magnified in (b). (**c,d**) EM image plane showing the blood vessel (B.v.) corresponding to the vessel shown in (ab) overlaid with the coarse affine transformation (see Fig. 3) of the LM-based axon reconstruction ax1_LM_^AT^; magnified in d. Note that ax1_LM_^AT^ does not match the ultrastructural features at high-resolution EM (d) due to the limited precision of ATBV. (**e,f**), EM plane overlaid with ax1 _LM_^AT^ (as shown in **c**) with bounding box for dense axon reconstruction (light blue) of edge length *λ*_align_ = 5 μm; demagnified in f, dashed rectangle denotes region shown in e. (**g**) Skeleton reconstructions of all n=193 ax1_EM_ candidates (light grey) traversing the search bounding box (as in e-f) (**h**) Reconstructed ax1_EM_ candidates (light grey) and the transformed fluorescent axon skeleton ax1_LM_^AT^ (black). Five candidate EM reconstructed axons with the most similar trajectories as ax1_LM_^AT^ highlighted in different colors. (**i**) Number of EM-reconstructed axons persistently within the λalign neighborhood of the LM-based template axon ax1_LM_^AT^. Note that at a distance of 43 μm from the seed center (e-h), only one possible ax1_EM_ candidate (green) remains. (**j**) Overlay of ax1_LM_^AT^ and the most likely ax1_EM_ candidate reconstruction. Axonal varicosities (filled circles) were independently annotated in LM and EM. Note the strong resemblance of the varicosity arrangements indicating that ax1_EM_ is the correctly matched EM equivalent of the LM signal (see Fig. S2 for statistical analysis). (**k**) EM plane with overlaid ax1_EM_ skeleton (dark green) and volume annotation of the corresponding EM axon ultrastructure (transparent green).

But how could we be sure that this EM axon was the one that had given rise to the fluorescence signal at the LM level? We first investigated the detailed morphology of this axon and identified axonal varicosities in the EM data. We then reconstructed the varicosities visible in the LM data and overlaid both (Fig. 4j). The similarity was visually striking (distance between LM and EM varicosities, 0.7 μm ± 0.6 (mean ± s.d., n=11); distance between LM and random EM varicosity distributions: 4.9 μm ± 2.1, (mean ± s.d., n=11 varicosities, n= 10^5^ draws, p<10^−5^, Randomization test, Fig. S2)). With this, we had found the EM axon that most likely explained the fluorescence signal reconstructed in the LM dataset (Fig. 4k).

We then continued to match other fluorescent axons given this first matched axon (Fig. 5). For this, we chose an axon in the LM data in proximity to the previously matched axon and selected an LM varicosity close to the axonal apposition as a seed point for the correspondence search in the EM data (Fig. 5a). Since each matched axon provided additional registration constraints beyond the initial blood vessel-based constraints, we could now use each matched axon to further refine an elastic free-form LM-to-EM transformation using the matched varicosities, branchpoints and other prominent structural features of the axons seen in both LM and EM (Fig. 4b,k). The iteratively refined registration together with the usage of varicosities as search criteria allowed us to substantially reduce the work load for matching subsequent fluorescent axons to their EM counterparts (Fig. 5b-e; in this example 34 axons were within a (2 × 2 × 3) μm^3^ search volume around the LM-varicosity, but only 6 of these had a varicosity). While the first match involved the reconstruction of 193 axons, the subsequent matches were successful after only several or even one reconstruction attempt and a few minutes of reconstruction time (Fig. 5e). The fact that all subsequent matches were successful (see varicosity correspondence as shown in Fig. 5f) implies that the previous matches were correct (since the probability of finding a correctly matched axon by chance after several chance matches is virtually zero). Using this strategy, we matched a total of 38 axons (27 in the red fluorescence channel, 11 in the green channel), using a total of 284 control points (7.5 ± 3.8 cps per axon (mean ± s.d), Fig. 5g).

**Figure 5.**
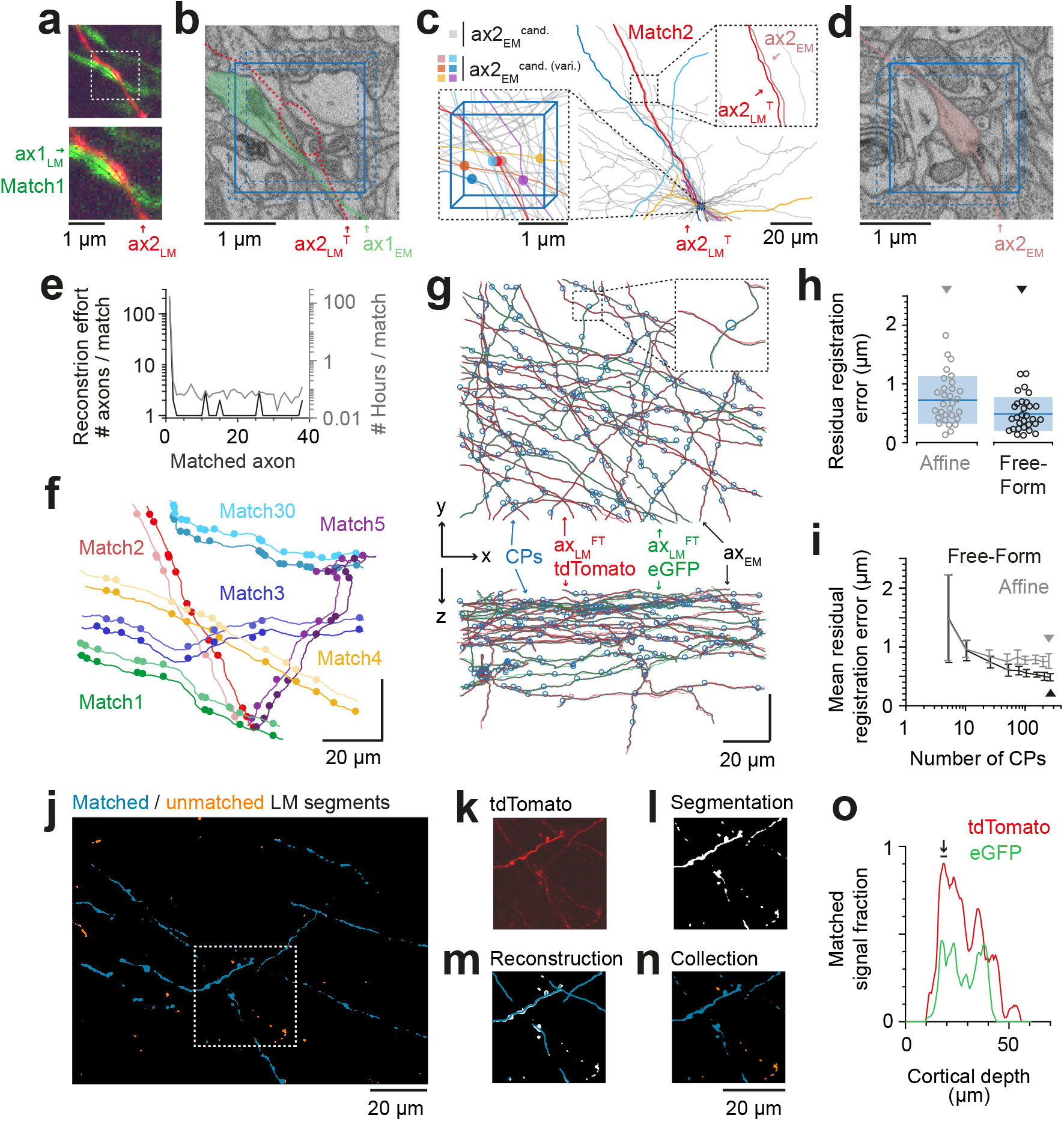
Iterative LM-to-EM axon matching. (**a**) Initial axon match ax1_LM_ and nearby axon ax2_LM_ with a varicosity in the proximity, both reconstructed in LM. (**b**) Matched axon ax1_EM_ (light green) and trajectory of axon 2 (ax2_LM_^T^) reconstructed at LM and transformed to EM (dashed red line) using a transformation T that was constrained using structural features of the initially matched ax1 (8 control points). (**c**) Search volume sized (2 × 2 × 3) μm^3^ (blue box) around the varicosity of ax2_LM_^T^ (see a-b) containing 34 candidate axons (gray) but only 6 with candidate varicosities (colored circles). One candidate axon had a similar trajectory as ax2LMT (right inset), identifying it as the correct match (see also varicosity pattern in f). (**d**) Varicosity in EM of axon ax2EM (red shading, see search template in b). (**e**) Reconstruction effort required to perform 38 iterative LM-to-EM axon matches (excluding control point placement). (**f**) Overlay of EM (light colors) and affine transformed LM (dark colors) axon reconstructions of 6 exemplary matched axon pairs with locations of axonal varicosities (independently reconstructed at LM and EM level, respectively). Note the similarity of axon trajectory and varicosity positions, also for the 30^th^ matched axon. EM axons were offset by 6 μm for visibility, see g for actual overlap quality. (**g**) Overlay of EM-reconstructed axon skeletons (black) with the matched LM-reconstructed skeletons transformed using a free-form transformation iteratively constrained by control points (CPs, blue circles) obtained from each consecutive axon match (shown transformation used a random subset of 250 (of 284) CPs). (**h**) Residual registration error of the match shown in g computed as the Euclidean distances ║ cp_EM_ – cp_LM_^FT^ ║ between n = 30 randomly picked CP pairs that had not been previously used to constrain the registration. Sample mean (blue line) and standard deviation (blue shading). (**i**) Average residual registration error (mean +− s.d., computed as in h) in dependence of the number of randomly chosen control points (CPs) used to constrain the transformation (n=10 bootstrapped CP sets, each). (**j-n**) Locally complete LM-to-EM matching of fluorescently labeled axons. One example image plane of LM dataset (at 18 μm depth) shown with EM-matched fluorescence signal fraction (blue) and yet unmatched segments (orange). To compute the matched fluorescence signal fraction, the raw LM data (k) was binarized and segmented (l, see Methods), overlaid with the EM-matched LM axon skeleton reconstructions (m) and only those segments overlapping with skeletons were counted as matched (n). (**o**) Matched signal fraction (in fraction of matched voxels) over dataset depth. For a range of 17-20 μm depth (black line, arrow), about 90% of the fluorescent voxels were explained by matched EM axons.

To quantify the improved registration precision obtained from the iteratively constrained LM-to-EM axon matches, we measured the residual registration error of the free-form registration (constrained by a random subset of 250 control points) on 30 other control points. The residual registration error was reduced to 487 ± 290 nm (mean ± s.d, Fig. 5h,i), thus about 6-fold smaller than in the coarse pre-alignment obtained before (cf. Fig. 3o).

To test whether we could perform a close-to-complete matching of LM-to-EM reconstructions for a certain depth range of the fluorescence dataset we matched all clearly visible fluorescently labelled axons at a depth of 17-20 μm in the red (tdTomato) channel (n=27 axons, total path length of 3.0 mm). We then computed a segmentation of the LM data and measured the fraction of the segmentation voxels overlapping with the EM-matched reconstructions (Fig. 5j-n). We found that in fact about 90% (86 ± 3%, mean ± s.d, n=7 LM image planes) of the fluorescent voxels were explained by the matched EM axons in this depth range (Fig. 5o).

In summary, investing a total of 181 work hours (178.4 hours for the first axon; 2.6 hours for the remaining axons) we obtained the EM equivalents of 38 fluorescently labeled axons in two fluorescence channels (total axonal path length of 4.6 mm). Since the matching of the first axon consumed most invested time, we checked whether by using the distribution of varicosities along EM and LM axons also for the initial match the time investment could be further reduced. In fact, when asking a second independent expert to perform the first LM-to-EM axonal match by also using varicosities as a search criterion, the invested time was only 7.5 minutes for the initial match. While this reduction in matching effort is well applicable to settings in which axonal varicosities can be detected, the more general but more laborious FluoEM approach of locally dense axonal reconstruction is expected to be applicable to other settings (for example transit axons that rarely form synapses). Together, we conclude that FluoEM is effective and efficient for LM-to-EM registration.

### Comparison of axon and synapse detection in LM and EM

The axons matched between the LM and EM volumes provided a dataset to calibrate in hindsight, how well axonal trajectories and branches can be reconstructed and how faithfully synapses can be detected in fluorescence LM data (Fig. 6).

**Figure 6.**
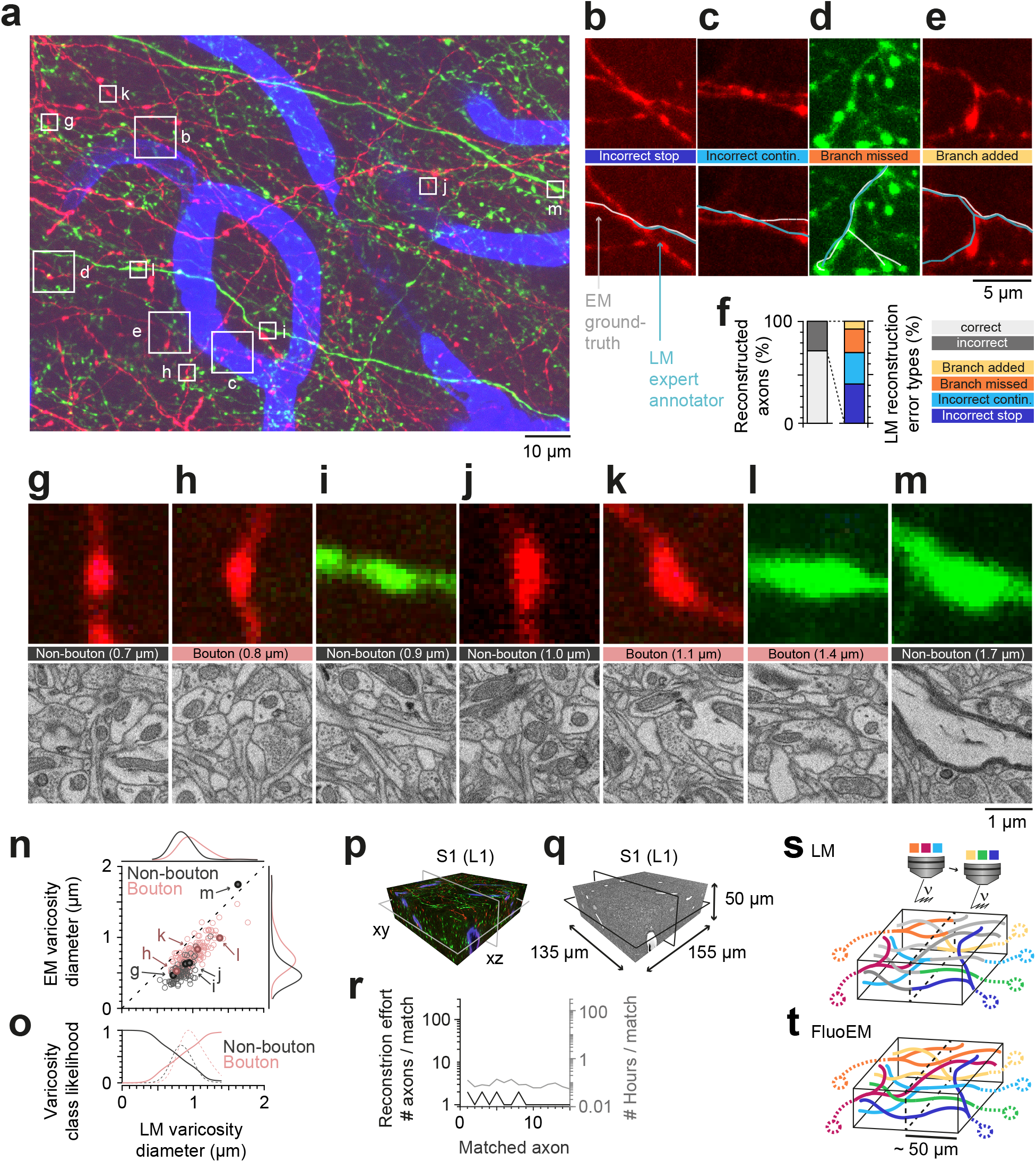
Calibration of axons and synapses in 3D fluorescent data using FluoEM, reproduction experiment, and color multiplexing in FluoEM. (**a**) Examples of axonal trajectories and axonal varicosities in 3D LM data (Maximum-intensity projection from 42 image planes at 17-35 μm depth; dataset and color code as in Fig. 3). (**b-f**) Post-hoc comparison of axonal reconstructions in LM data and their EM counterparts as identified by FluoEM, which were considered ground truth. Examples of LM reconstruction errors (b-e) and their prevalence (f). Note that while about 70% of axons were correctly reconstructed at the LM level, for about 30%, reconstruction errors such as missed branches or incorrect continuations occurred. (**g-m**) Validation of presynaptic bouton detection in fluorescence data: Examples of axonal varicosities imaged using LM (top row) and EM (bottom row). EM was used to determine synaptic (bouton) vs. non-synaptic varicosities. Size of varicosity reported as sphere-equivalent diameter (see Methods). (**n**) Relation between the sphere equivalent diameter of axonal varicosities imaged in EM vs. LM for synaptic (red, bouton) and non-synaptic (black) varicosities. (**o**) Likelihood of varicosities to be synaptic (red) vs. non-synaptic (black) over LM varicosity sphere equivalent diameter, computed from the respective (prevalence-weighted, kernel density estimated) probability densities (dashed, see Methods). (**p-r**) Second FluoEM experiment from upper cortical L1 in S1 cortex (p: LM dataset subvolume corresponding to (q); q: EM high-res dataset with dimensions) and reconstruction effort for LM-to-EM matching of axons (r). (**s,t**) Color multiplexing strategy to increase color space without increasing photon dose in FluoEM. If labeled axons traverse a large volume, then 3D LM imaging can be split into subvolumes of minimal dimension about 50 μm (given by d_recon_ and d_unique_, see Fig. 2) (s), and axon identity can be propagated into the remaining volume via EM reconstruction (t) using the FluoEM approach.

We first asked a different expert annotator to reconstruct the previously matched axons in the LM dataset, this time without an explicit bias to stop reconstruction at uncertain locations (cf. Fig. 2c), and without knowledge of the matched EM axon’s shape or trajectory. We then compared these axonal reconstructions performed in the LM dataset with the corresponding EM trajectories of the axons, which we considered the ground truth for the detection of branchpoints, the determination of the axonal morphology and the presence of presynaptic boutons (Fig. 6a-f). We found that while a large fraction of axons were reconstructed faithfully at the LM level (27 of 38 axons, 71%), in 29% of cases the LM reconstruction contained an error (Fig. 6f). The largest fraction of errors was related to incorrectly terminated reconstructions (“split”, 38%, Fig. 6b,f), followed by an incorrect continuation of the LM-reconstructed axon at locations where multiple fluorescent axons overlapped (31% of errors, Fig. 6c,f). Furthermore, missed (23%) and added (8%) branches occured (Fig. 6d,e,f). Together, this data confirmed that already at intermediate labeling density, fluorescence-based axon reconstruction in cortex is error-prone.

We then identified 255 axonal varicosities in the LM data and investigated their ultrastructure at the EM level based on the successfully matched axonal counterparts (Fig. 6a,g-o). As the examples indicate (Fig. 6g-m), many varicosities in the LM in fact correspond to presynaptic boutons (63%, n=161 of 255, Fig. 6h,k,l). A substantial fraction of LM-varicosities, however, are without evidence for a presynaptic specialization and instead contain only mitochondria (n=54, Fig. 6g,j), or are hollow thickenings (n=39, Fig. 6i) or myelinated (n=1, Fig. 6m). Notably, while the inference of synaptic boutons from LM varicosities was dependent on the size of the LM varicosity (Fig. 6n) with larger varicosities more likely synaptic, there was no LM varicosity size range that allowed unequivocal inference of a synapse from the LM morphology (Fig. 6n,o).

### Reproduction experiment

Finally, in order to assure the reproducibility of FluoEM, we conducted a second correlated imaging experiment in layer 1 of mouse S1 cortex (LM dataset sized (1027 × 820 × 84) μm^3^, EM dataset sized (155 × 135 × 50) μm^3^, Fig. 6p-q), in which we successfully matched 15 axons between LM and EM as a proof of principle (matching effort 1.3 hours total, 5.3 ± 1.6 minutes per axon, Fig. 6r).

## DISCUSSION

We report FluoEM, a set of experimental and computational tools for the virtual labeling of multiple axonal projection sources in connectomic 3D EM data of mammalian nervous tissue. At its core is the notion that dense EM circuit reconstruction allows to reformulate the alignment between data obtained at micrometer resolution (LM) to data obtained at nanometer resolution (EM) to the best-matching between a sparse subset of labeled axons (LM) and all possible axons (EM). We find that in mammalian neocortex, densely reconstructed axons have unique trajectories on a spatial scale that is compatible with currently achievable 3D EM dataset sizes, allowing the FluoEM concept to be successfully applied. With this, FluoEM allows the enriching of EM-based connectomes in mammalian neuropil with the large range of multi-color encoded information already successfully expressed in axons at the light-microscopic level.

One key prerequisite for FluoEM is the uniqueness of axonal trajectories in dense neuropil. We have so far determined the uniqueness length for several layers of the mouse cerebral cortex (Fig. 2), where this length is consistently below 40 μm (90^th^ percentile, see Fig. 2), thus in a range that can be faithfully reconstructed at the LM level (Fig. 2a-c). For a broad range of cortical and subcortical nervous tissue (basal ganglia, thalamus, hypothalamus, nuclei of the brain stem etc.) in which locally dense circuits coexist with incoming projection axons from distant sources, FluoEM can be applied on the spatial scales described here. Cases in which dense neuropil contains highly anisotropic axons (for example axon bundles in the white matter of mammalian brains) will likely imply a longer scale on which axonal trajectories become unique, and therefore require longer reconstruction lengths in the fluorescence data.

We have developed FluoEM using a particular 3D EM imaging technique, SBEM. Since FluoEM only relies on the underlying geometry of axonal trajectories, which we have shown to be unique on scales achievable with all current 3D EM techniques (Fig. 2g-k), FluoEM is not restricted to the combination with SBEM. Rather, other 3D EM imaging approaches, especially FIB-SEM (Heymann, Hayles et al. 2006, Knott, Marchman et al. 2008, Hayworth, Xu et al. 2015, Xu, Hayworth et al. 2017) and ATUM-SEM (Schalek, Kasthuri et al. 2011, Kasthuri, Hayworth et al. 2015) that provide faithful axonal reconstructions over at least about 40-50 μm extent in 3 dimensions will be useable for the presented approach.

While FluoEM in principle allows the identification of as many axonal projection sources in a single connectomic experiment as can be encoded at the light-microscopic level, it is practically limited by the number of fluorescence channels that can be acquired in sequence without exerting damage to the tissue. In our experiments, tissue exposed to 2 hours of continuous LM imaging with acquisition of 3 parallel fluorescence channels was still well usable for subsequent 3D EM circuit reconstruction. To acquire more fluorescence channels without increasing the local photon dose in the tissue, one can exploit the fact that in FluoEM, it is not necessary to image the axons of interest in their entirety at the LM level. Rather, it is sufficient to LM-image volumes of 40-50 μm in extent, as long as all axons of interest pass through that volume, and to reconstruct the complete axons in EM. Therefore, a setting in which the volume of interest is split into several subvolumes, in each of which a limited number of fluorescence channels is acquired, could multiplex the color range of FluoEM (Fig.6s-t) without increasing the exposure time in the LM step.

In LM imaging of mammalian nervous tissue, labeling density of axons poses a notorious limitation as long as axonal trajectories were to be reconstructed from the LM data (labeling more than about 1 in 1000 to 10,000 axons impedes reconstruction (Helmstaedter 2013)). With FluoEM, labeling density is only limited by the need to follow axons over the uniqueness length (Figs. 2,3). Since this depends on the overall alignment precision (Fig. 2k), a setting in which one fluorescence channel contains sparsely labeled axons to provide an accurate pre-alignement (see Fig. 5g,h), and the other fluorescence channels are densely labeled is a possible solution, in which LM-reconstructable stretches of axons as short as 5 μm would be sufficient for co-registration (Figs. 2k and 5h).

Finally, the reconstruction work load in FluoEM requires a quantitative discussion. In settings where axonal varicosities can be used as additional constraints on the matching (Fig. 5a-d), the LM-to-EM match consumed about 10-20 minutes per axon (including the placement of control points). With this, a full matching of a sparsely stained volume sized (1 × 1 × 0.1) mm^3^, for example a large area of layer 1 in the mouse neocortex, would thus likely consume about 2700 work hours in FluoEM (using our efficient viewer webKnossos (Boergens, Berning et al. 2017)). While this is a considerable time investment, it favorably compares to other resource investments in biomedical imaging. Furthermore, with the development of increasingly reliable automated approaches for the reconstruction of axons and synapses at the EM level (Berning, Boergens et al. 2015, Staffler, Berning et al. 2017), a further speedup of FluoEM is conceivable in which partially automated matching contributes. In mammalian circuits where axons may not show local diameter variations that can be used for co-registration, the FluoEM approach using dense axonal reconstruction (Fig. 2) is applicable.

Together, the presented methods overcome the major hurdle of limited color space for identifying multiple axonal input sources in parallel in large-scale and dense electron microscopic reconstruction of mammalian neuronal circuits.

### Author contributions

Conceived and supervised the project: MH. Performed experiments and analyses: FD. Contributed reconstructions: AK. Contributed EM data: KMB. Wrote the paper: MH and FD with contributions by all authors.

## Acknowledgements

We thank Kevin Briggman and Gilles Laurent for discussions, Marcel Beining and Kun Song for comments on the manuscript, Alessandro Motta, Manuel Berning, Benedikt Staffler and Emmanuel Klinger for computational advice, Stephan Junek for support with light microscopic imaging, Johannes Letzkus for advice and support with viral injections, Heiko Wissler and Dalila Rustemovic for tracer management. We thank Matt Jacobson for providing the “Absolute Orientation – Horn’s method” Matlab central package we used in our affine registration workflow. We thank Dirk-Jan Kroon for providing the “B-spline Grid, image and point registration” Matlab central package we used for our free-form registration workflow. We thank Aymun Al-Shaboti, Mahmoud Aly, Carolin Arras, Nagihan Aydin, Susanne Babl, Danny Baltissen, Anne Bamberg, Feron Basoeki, Natalie Berghaus, Alfred Berghoff, Lisa Bezzenberger, Nicola Böffinger, Svenja Bohne, Alexander Brandt, Oliver Brandt, Julius Buss, Lars Buxmann, Deniz Celik, Hanaa Charif, Nadia Cipta, Linda Decker, Kristina Desch, Tim Engelmann, Theresa Ernst, Jordi Espino Martinez, Benjamin Fani Sani, Theresa Fueller, Daniel Goffitzer, Victoria Gosch, Jennifer Hartel, Hanna Hees, Björn Heftrich, Julian Heller, Robert Hülse, Oana Ilea, Raphael Jakob, Raphael Jakoby, Alexander Jost Lopez, Marcel Jüngling, Vanessa Kalbert, Mehmet Karabel, Lennart Kirchner, Raphael Kneißl, Plamen Kondev, Patricia König, Katharina Kramer, Franziska Krämer, Leonhard Kreppner, Marc Kronawitter, Jülide Kubat, Benjamin Kuhl, Derya Kurt, Eike Laube, Maria Leondaraki, Rebecca Lotz, Carina Lossnitzer, Luisa Lutz, Saskia Mehlmann, Isabell Metz, Nils Plath, Anna-Lena Possmayer, Leonard Präve, Maximilian Präve, Sylvia Reibeling, Sascha Reichel, Anna Rix, Claudio Sabatelli, Clemens Schumm, Lilli Schütz, Britta Stiehl, Kira Trares, Simon Umbach, Marco Werr, Jannik Winkelmeier, Timm Winkelmeier and Susanne Zimbelmann for neurite reconstruction.

## METHODS

### Animal Experiments

All experimental procedures were performed according to the law of animal experimentation issued by the German Federal Government under the supervision of local ethics committees and according to the guidelines of the Max Planck Society.

### Virus Injection

Young adult (p28) male wild-type mice (C57BL/6j) were anesthetized with isoflurane (induction: 4%, maintenance: 2%) in medical oxygen and fixed in a stereotaxic frame (Model 1900, Kopf Instruments, USA). The core body temperature was maintained at 37°C using a feedback-controlled heating pad (DC Temperature Control System, FHC, USA). Systemic analgesia was provided by subcutaneous injection of 2 mg/kg Meloxicam (Metacam, Boeringer-Ingelheim, Germany) and 100 mg/kg Metamizol (Metamizol WDT, WDT, Germany) prior to surgery. Local anesthesia was provided by injecting 16.7 mg/kg Ropivacaine (Naropin, AstraZeneca, Switzerland) under the scalp. To anterogradely label projections originating from M1 and S2 cortex, adeno-associated viruses expressing eGFP (AAV1.CAG.FLEX.EGFP, 2.4×10^13^ GC/ml, Penn Vector Core, USA) and tdTomato (AAV1.CAG.FLEX.tdTomato, 1.5×10^13^ GC/ml, Penn Vector Core, USA) were injected using freshly pulled 1.5 mm borosilicate glass capillaries at the following coordinates: 1.0 mm anterior of bregma, 1.0 mm lateral of the midline, 0.7 mm below the cortical surface (M1) and 0.6 mm posterior of bregma, 4.1 mm lateral of the midline, 1.3 mm below the cortical surface (S2). Both AAV.FP solutions were mixed in a 2:1 ratio with AAV. Cre solution (AAV1.CamKII0.4.Cre, 4.8×10^13^ GC/ml, Penn Vector Core, USA) prior to injection. At each site approximately 50 nl were injected using a pressure injection system (PDES, NPI, Germany).

### Transcardial Perfusion

Three weeks after virus injection, mice were subcutaneously injected with 0.1 mg/kg Buprenorphine (Buprenovet, Recipharm, France) and 100 mg/kg Metamizol. General aneasthesia was introduced with 4% isoflurane and maintained with 3% isoflurane in medical oxygen. Mice were transcardially perfused with 10 - 15 ml flushing solution containing a lipophilic far-red fluorescent dye for blood vessel labeling (150 mM Sodium Cacodylate, 5 μM 1,1’-Dioctadecyl-3,3,3’,3’-Tetramethylindodicarbocyanine (DiD, Sigma-Aldrich, USA), pH 7.4) followed by 50 - 60 ml fixative solution (80 mM Sodium Cacodylate, 2.5% Paraformaldehyde, 1.25% Glutaraldehyde, 0.5% CaCl_2_, pH 7.4) using a syringe pump (PHD Ultra, Haravard Apparatus, USA) at a flow rate of 10 ml / min. After perfusion, mice were decapitated and the connective tissue around the skull was removed. The head was immersed in fixative solution for approximately 24 hours at 4°C.

### Sampling

After extracting the brain from the skull and placing it in a dish filled with storage buffer (150 mM Sodium Cacodylate, pH 7.4) coronal vibratome (HM650V, Thermo Fisher Scientific, USA) sections of 100 – 150 μm thickness were made until the target position along the rostro-caudal axis was reached at 1.6 mm posterior of bregma. A coronal slab of 1 mm thickness was then cut off the remaining brain from which a cylindrical sample containing cortical layers 1 – 5 was extracted from 1.3 mm lateral of the midline using a medical 1.5 mm biopsy punch (Integra, Miltex, USA).

### Confocal Imaging

Prior to confocal imaging, the fixed brain tissue sample was washed for 30 minutes in storage buffer containing a mild reducing agent (150 mM Sodium Cacodylate, 2 mM reduced Glutathion, pH 7.4) and then transferred into a storage buffer-filled imaging chamber (Cover Well, Grace Bio-Labs, USA). Using an inverted Confocal Laser Scanning Microscope (LSM 880, Zeiss, Germany) and a 40x water objective (C-Apochromat 40x/1.2 W Korr FCS M27, Zeiss, Germany), a 3-channel (eGFP labeled axons, tdTomato labeled axons, DiD labeled blood vessels) image stack of dimensions (828 × 828 × 75) μm^3^ was acquired at a voxel size of (104 × 104 × 444) nm^3^ in a single scanning track.

### EM Staining

Immediately after imaging the sample was removed from the imaging chamber and immersed in 2% OsO_4_ aqueous solution (Serva, Germany) in cacodylate buffer (150 mM, pH 7.4). After 90 minutes the solution was replaced by 2.5% ferrocyanide (Sigma-Aldrich, USA) in cacodylate buffer (150 mM, pH 7.4) and incubated for another 90 minutes. The solution was subsequently exchanged for 2% OsO_4_ in cacodylate buffer (150 mM, ph 7.4) and incubated for another 45 minutes. The sample was then washed first for 30 minutes in cacodylate buffer and then for another 30 minutes in ultrapure water. Next, the sample was incubated in a 1% thiocarbohydrazide (Sigma-Aldrich, USA) at 40°C for 45 minutes and washed twice in ultrapure water for 30 minutes each before immersing it 2% unbuffered OsO_4_ aqueous solution. After 90 minutes the sample was again washed twice for 30 minutes each in ultrapure water and subsequently immersed in 1% uranyl acetate (Serva, Germany) aqueous solution at 4°C overnight. On the next day the sample (still immersed in uranyl acetate solution) was heated to 50°C in an oven (Universal Oven Um, Memmert, Germany) for 120 minutes. After two washing steps with ultrapure water lasting 30 minutes each, the sample was incubated in a 20 mM lead aspartate (Sigma-Aldrich, USA) solution (adjusted to pH 5 with 1N KOH) at 50°C for 120 minutes. The sample was then washed twice in ultrapure water again for 30 minutes each. Unless specified otherwise all steps were carried out at room temperature.

### EM Embedding

After concluding the staining procedure the sample was dehydrated by immersing it for 30 minutes each at 4°C in a graded ethanol (Serva, Germany) series of 50% ethanol in ultrapure water, 75% ethanol in ultrapure water and 100% ethanol, respectively. The sample was then immersed three times in acetone (Serva, Germany) 30 minutes each at room temperature. After completing the dehydration steps, the sample was infiltrated with a 1:1 mixture of acetone and Spurr’s resin (Sigma Aldrich, USA) with a component ratio of 4.1 g ERL 4221, 0.95 g DER 736 and 5.9 g NSA and 1% DMAE at room temperature for 12 hours on a 45 degree rotator in open 2 ml reaction tubes (Eppendorf, Germany). Infiltrated samples were then incubated in pure resin for 6 hours, placed in embedding molds and cured in a preheated oven at 70°C for 48 – 72 hours.

### EM stack acquisition

The embedded sample was trimmed with a diamond-head milling machine (EM Trim, Leica, Germany) to a cube of approximately (1.5 × 1 × 1) mm^3^ dimension and mounted on an aluminium stub with epoxy glue (Uhu Plus Schnellfest, Uhu, Germany) in tangential orientation, so that the pia mater was located at the top of the block face. The thin layer of resin covering pia was carefully removed with a diamond-knife ultramicrotome (UC7, Leica, Germany). The smoothed tissue block was then gold-coated (ACE600, Leica, Germany) and placed in a custom-built SBEM microtome (courtesy of W. Denk) mounted inside the chamber of a scanning electron microscope (FEI Verios, Thermo Fisher Scientific, USA). Microtome movements and image acquisition were controlled using custom-written software packages interfacing with motor controllers and the microscope software’s API. A low resolution image stack of dimensions (1500 × 1000 × 10) μm^3^ was acquired at a pixel size of (270 × 270) nm^2^ and a cutting thickness of 200 nm. After correspondences with the confocal image stack were successfully identified, a high resolution image stack sized (130 × 110 × 85) μm^3^ was acquired at a pixel size of (11.24 × 11.24) nm^2^ and a cutting thickness of 30 nm (Fig. 3).

### Image post-processing

The LM image stack (Fig. 2a-c, Fig. 3) consisted of 16 × 153 (xy x z) image tiles (2048 × 2048 pixels each) acquired at 8 bit color depth for each of the 3 color channels, resulting in 29 GB of uncompressed data. The 2448 image tiles were stitched into a global reference frame using the Zen software package (Zeiss, Germany). The LM image volume was then rotated so that the orientation of the superficial blood vessel pattern coarsely matched the orientation of the blood vessel pattern visible in the EM overview images. The images were then converted into the Knossos (Helmstaedter, Briggman et al. 2011) format and transferred to a data-store hosted at the Max Planck Compute Center in Garching from which they can be accessed via our online data viewer webKnossos ((Boergens, Berning et al. 2017), webknossos.org).

The EM image stack acquired was comprised of 20 × 2768 (xy x z) image tiles (3072 × 2048 pixels each) with 8 bit color depth, resulting in 326 GB of uncompressed data. The 55360 resulting tiles were stitched into a global reference frame using custom written Matlab software based on the approach described in (Briggman, Helmstaedter et al. 2011). However, instead of cross-correlation we used SURF feature detection to estimate relative shift vectors and their confidence. We then used a weighted least-squares approach to retrieve a robust globally optimal solution given these parameters. The EM data was then also converted into the Knossos format and transferred to a data-store hosted at the Max Planck Compute Center in Garching from which they can be accessed via our online data viewer webKnossos.

### Measurement of LM axon reconstructable pathlength (d_recon_)

For estimating the distribution of the reconstructable pathlength *d_recon_* in the LM data, a (40 × 40 × 48) μm^3^ bounding box was placed in a subvolume of the dataset with representative labeling density and all visible axons traversing this bounding box were skeleton-reconstructed independently by two expert annotators using webKnossos. Then the path-lengths of the resulting skeleton representations were measured by calculating the cumulative sum of the Euclidian distances between all connected nodes and a histogram over the distances was computed (Fig. 2c).

### Measurement of EM axon uniqueness pathlength (d_unique_)

To measure the uniqueness distance *d_unique_* of cortical axons in different cortical depths, a rectangular bounding box of edge length *λ_align_* was placed at the center of each dataset and all axons traversing this bounding box were reconstructed, resulting in a set of axon skeletons. To measure *d_unique_* for a given axon *ax_i_* a marching sphere with radius *r* ≈ *λ_align_* was placed iteratively around each node 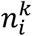 and all neighboring axons *ax_j_* with at least one of their nodes 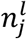 residing within the sphere so that 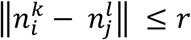 were identified, resulting in a binary neighbor vector indicating the presence (*true*) or absence (*false*) of each putative neighboring axon at each sphere (node) location. The results were then sorted according to the Euclidian distance between the respective sphere (node) location and the bounding box origin. Whenever a value of the distance sorted neighbor vector changed from true to false at consecutive distances *d* and *d* + 1 the respective distance *d* was considered the dropout distance for a given neighbor axon *ax_j_*. We define *d_unique_* and 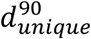 as the Euclidean distances from the origin for which all or 90% neighboring axons have dropped out, respectively (Fig. 2f,g-i).

### Coarse LM-EM alignment

Based on the characteristic pattern of large superficial blood vessels visible both in the LM data and the low-resolution EM data (Fig. 3d,e) a coarse correspondence was visually established between the two datasets. An approximate bounding box was applied to the LM dataset defining the sub-volume corresponding to the acquired high resolution EM dataset. Subsequently all blood vessels were traced using webKnossos. Based on characteristic blood vessel bifurcations, 7 pairs of blood vessel skeleton nodes were defined as corresponding control point pairs *CP* = (*CP_EM_,CP_LM_*) with 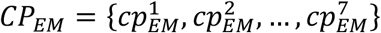 and 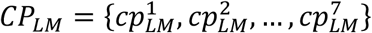 used to constrain a three dimensional affine transformation *AT_BV_* (Fig. 3h-o).

### Affine registration

Given a set of control point pairs and an initial scaling vector resulting from the ratio between the nominal LM and EM voxel sizes we computed an initial affine transformation using Horn’s quaternion-based method (Horn 1987). We then iteratively optimized the scaling vector by minimizing the least squares residual registration error. We then applied the resulting transformation to the nodes of skeleton reconstructed LM blood vessels or axons to obtain affine transformed skeletons registered to their corresponding structures in EM reference space.

### Free-from registration

Given a set of control point pairs and an initial scaling vector we first computed the optimal affine registration as described above. We then used a b-spline interpolator based approach (Lee, Wolberg et al. 1997, Rueckert, Sonoda et al. 1999) to compute an optimal free-form registration given a set of control points, an initial b-spline knot spacing and a number of bisecting mesh refinement steps. We found a combination of 32 μm initial spacing and 4 refinement steps to give us an optimal compromise between registration precision and robustness. We then applied the resulting deformation grid to the nodes of skeleton reconstructed LM axons to obtain free-form transformed skeletons registered to their corresponding structures in EM reference space.

### Initial axon matching

As a candidate for the initial axon matching an axon *ax*1_*LM*_ in spatial vicinity of a blood vessel bifurcation was picked and reconstructed in the LM dataset using webKnossos. The previously constrained affine transformation was then used to transform *ax*1_*LM*_ into the EM reference space so that 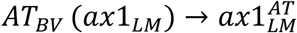. Because *ax*1_*LM*_ was chosen close to one of the corresponding blood vessel bifurcations, it was possible to estimate the local affine transformation error by applying the same transformation to the blood vessel skeletons and measuring the offset at the closest bifurcation. Based on this offset a local translative correction 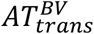 was defined and used to locally correct the offset of the first axon candidate: 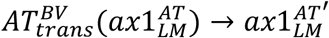. A bounding box was then defined around a node 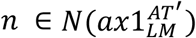 close to the blood vessel bifurcation and all axons traversing this bounding box were reconstructed. The true corresponding axon *ax*1_*EM*_ amongst all of these possible axon candidates was selected by the marching sphere method (see methods: measurement of EM axon uniqueness pathlength, Fig. 2, Fig. 4i) followed by a validation based on varicosity patterns (Fig. 4j).

### Iterative axon matching and free-form transformation

After successfully matching a given axon we used its varicosities and branchpoints as additional constraints to update our registration, thereby increasing local registration precision substantially. By applying these updates and carefully choosing an axon in the spatial vicinity of a previously matched as the next matching candidate we were able to reduce the search volume from (5 μm)^3^ to (2-3 μm)^3^ depending on the distance of the seed point on the candidate axon to the closest previously matched constraint and the general registration precision in that area. Furthermore, by using a varicosity as a seed point around which the search volume would be centered we were able to even further reduce the number potential EM candidates to only such axons displaying such a structural feature within the reduced search volume (Fig. 5a-d). Only if we did not succeed finding the match within the small bounding box, its volume was incrementally increased until the match was found. Using this method, we were able to identify the match for every axon we attempted.

### Registration error measurements

The residual registration error of the blood vessel-based affine registration was measured by applying the affine transformation *AT^BV^* to the nodes of the LM blood vessel skeleton representations and measuring the Euclidean distances between the EM control points *CP_EM_* and the transformed LM control points 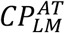 located on the respective skeletons. In this case we used the same *n*(*CP_residual_*) = 7 control point pairs to constrain the transformation and measure the residuals (Fig. 3o).

The residual registration error of the axon based affine registration was measured by applying the affine transformation *AT* to the nodes of the LM axon skeleton representations and measuring the Euclidean distances between the EM control points *CP_EM_* and the transformed LM control points 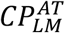 located on the respective skeletons. We randomly picked *n*(*CP_residual_*) = 30 control point pairs, none of which had been used before to constrain the transformation, to measure the residuals (Fig. 5h).

The residual registration error of the axon based free-form registration was measured by successively applying the affine transformation *AT* followed by the free-form transformation *FT* to the nodes of the LM axon skeleton representations and measuring the Euclidean distances between the EM control points *CP_EM_* and the transformed LM control points 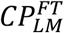 located on the respective skeletons. We randomly picked *n*(*CP_residual_*) = 30 control point pairs, none of which had been used before to constrain the transformation, to measure the residuals (Fig. 5h).

To measure the effect the number of control points exerts on the residual registration error we randomly picked *n*(*CP_constrain_*) = [5, 10, 25, 50, 75, 100, 150, 200, 250] control points from a population of *n*(*CP_total_*) = 284 control points and computed the registration. We then randomly picked an additional *n*(*CP_residual_*) = 30 control points none of which were used in the step before for constraining the registration. We repeated this procedure 10 times for each number of control point pairs and reported the average and standard deviation of the respective sample residual mean (Fig. 5i).

### Varicosity diameter measurements

We define axon varicosities as roughly ellipsoid-shaped local diameter increases for EM and as roughly ellipsoid-shaped local signal width and/or intensity increases for LM. The equivalent sphere diameters *d* reported for the varicosities were calculated based on measured volumes *V* using 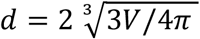. EM varicosity volumes *V_EM_* were measured by first computing an over-segmentation using SegEM (Berning, Boergens et al. 2015), manually collecting segments belonging to a given varicosity and summing the number of voxels multiplied by the respective voxel volume. LM varicosity volumes were obtained by manually annotating the three principal axes of the ellipsoid-shaped varicosities, extracting the intensity profiles along these axes and fitting Gaussian functions to them. Finally, *V_LM_* was estimated as the ellipsoid volume resulting from the three full-width half-maximums of these Gaussians relative to the local baseline intensity (Fig. 6g-n).

### Varicosity class likelihood estimation

To estimate the varicosity class likelihood we performed a kernel density estimation (bandwidth 100 nm) and weighted the estimated probability densities with the relative prevalence of the bouton vs non-bouton class (Fig. 6o, dashed lines). To obtain the class likelihood we divided the respective class probability density by the sum of both probability densities (Fig. 6o, solid lines).

### Comparison of axon detection in LM and EM

To test the reliability of axon reconstruction in LM we asked an expert annotator to again reconstruct all 38 axons that had been successfully matched before (by another expert), without knowledge of their shape or trajectories. To initialize these reconstructions, we randomly selected a central skeleton node and its two direct neighbors from each axon (only nodes of node degree 2 were considered). The expert annotator was then instructed to complete the LM axon reconstructions based on these initial locations using webKnossos; the expert was asked to reconstruct faithfully but also to attempt a full axonal reconstruction (thus relieving the bias towards split reconstructions applied in the calibrations of Fig. 2c). The resulting reconstructions were then compared to the EM skeletons by visual inspection, and errors counted (Fig. 6a-f).

### FluoEM for larger volumes: Extrapolation

In a (130 × 110 × 70) μm^3^ volume we matched 38 axons constituting approximately 36% of the total fluorescence signal in both color channels. Assuming an average matching time effort of 5 minutes per axon and an average control point placement time of 10 minutes per axon, a total time effort of 9.5 hours would be required to match the volume with 36% coverage. To achieve 100% coverage for the matched volume, 9.5h x (100% / 36%) ≈ 27h would be necessary. Thus, in order to match a volume of (1000 × 1000 × 100) μm^3^ one can estimate the time effort to be approximately 27h x (1000μm /130μm) x (1000μm /110μm) x (70μm /100μm) ≈ 2700h.

### Statistical tests

#### All statistical tests were performed using MATLAB

To test whether the observed differences in 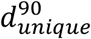(Fig. 2j,k) between L1, L2/3 and L4 are caused by the varying numbers of traversing axons encompassed in the (5 μm)^3^ bounding boxes (L1: n=220 axons, L2/3: n=207 axons, L4: n=128 axons), we repeatedly (n=30 draws) bootstrapped random subsamples of 128 axons from the L1 and L2/3 bounding box axon samples (without replacement) and measured 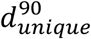. In both cases (L1: p=0.54, L2/3: p=0.14) the bootstrapped axon sets were indistinguishable from 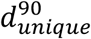 of the L4 bounding box axon sample (one sample t-tests, see Fig. S1).

To validate the initial axon match we compared the similarity of the measured varicosity distributions on the LM axon and EM candidate axon (Fig. 4j). To determine the probability of finding a matching varicosity pattern by chance we performed a randomization test in which we repeatedly (n=100,000 draws) assigned the measured number of EM varicosities (n=11) to random positions (nodes) on the EM skeleton. We then calculated the mean Euclidian distance between the measured varicosity positions on the affine transformed LM axon and the (either measured or randomly drawn) positions on the EM axon and found it is unlikely (p=10^−5^) to find a match as good or better as the match based on the actually measured varicosity positions (Fig. S2).

---

**Figure S1.**
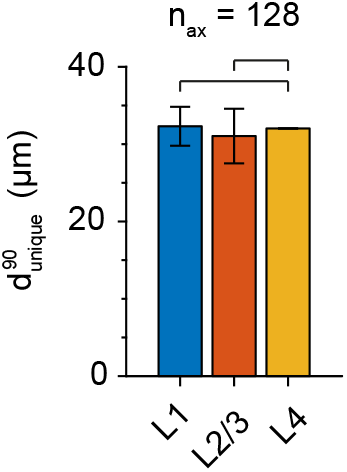
Axonal uniqueness length *d_unique_* for different cortical layers corrected for numbers of contained axons. Bars indicate mean 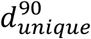 (error bars: s.d.) of the bootstrapped L1 and L2/3 axon bounding box samples (n=128 out of L1: n=220, L2/3: n=207 axons per random subsample, n=30 draws). For L4 (n=128 axons contained in bounding box sample) bar indicates 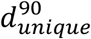. Statistical tests do not indicate a significant axon uniqueness length difference between the bootstrapped L1 and L2/3 distributions and the L4 sample mean (L1: p=0.54; L2/3: p=0.14, see Methods: Statistical tests).

**Figure S2.**
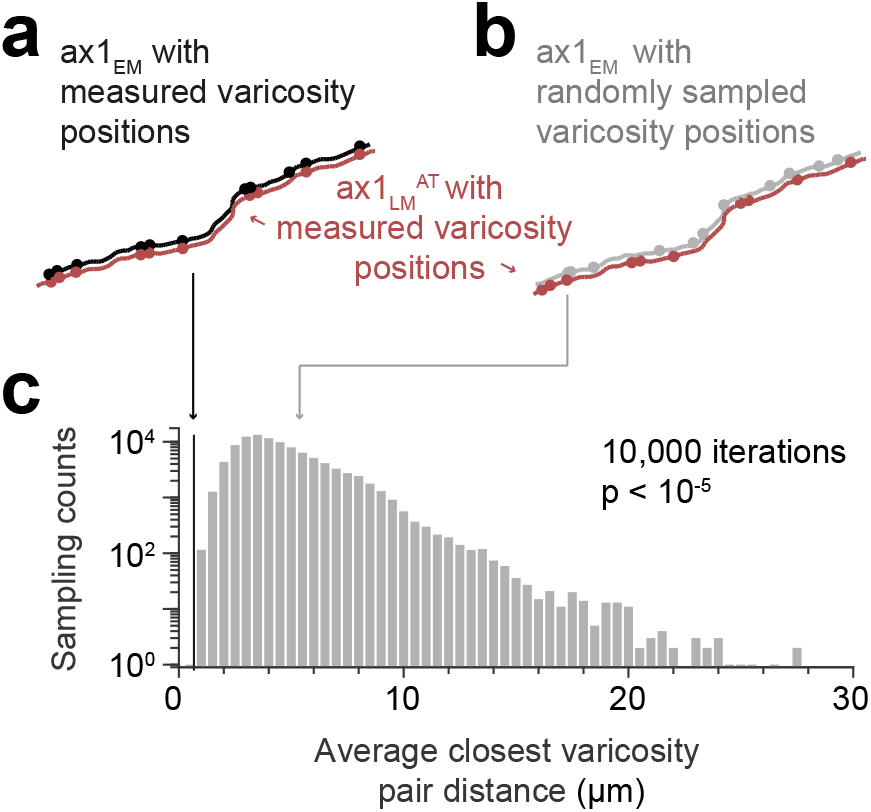
Similarity of matched varicosity patterns between LM and EM. (**a**) Independently measured varicosities (red and black circles) highlighted on the skeleton overlay of the initial affine transformed LM axon (ax1_LM_^AT^, red) and the EM match candidate (ax1_EM_, black) (compare Fig. 4j). (**b**) Overlay as in (a) but with ax1_EM_ varicosity positions (grey circles) randomly sampled. (**c**) Distribution of the average varicosity pair distance between the measured varicosities on ax1_LM_^AT^ and the respective closest available varicosity on ax1_EM_ for the measured (black line) and randomly sampled (grey histogram bars) positions. The probability of randomly picking (n=10,000 draws) varicosity positions with an average bouton pair distance as small as or smaller than the measured positions (black line, d = 0.67 μm) is p ≤ 10^−5^ (see Methods: Statistical tests).

**Figure S3.**
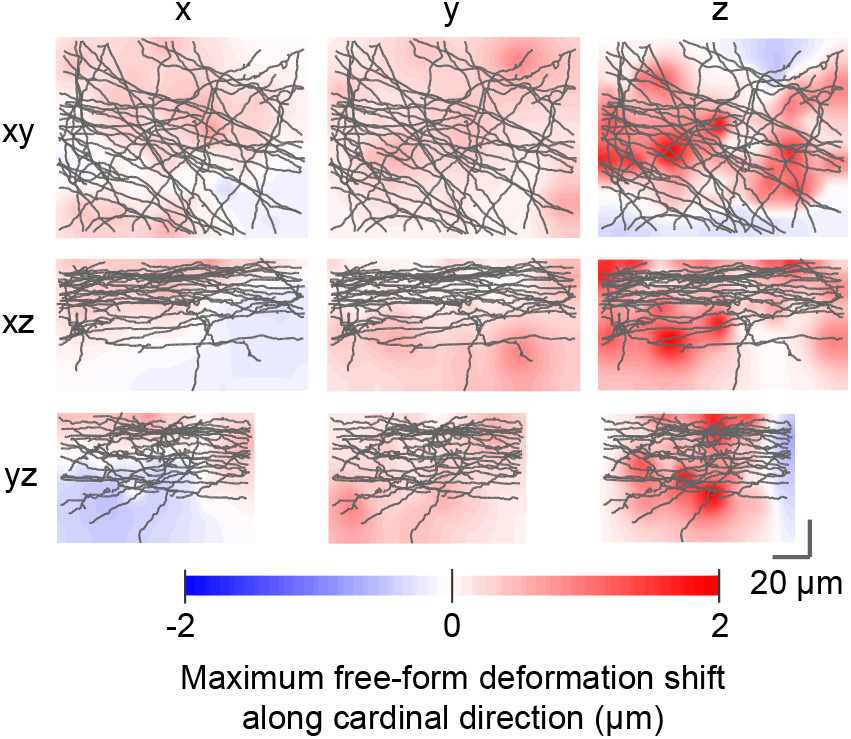
Deformations of axons in LM vs. EM data. The relative Euclidean displacements resulting from the free-form registration relative to the affine registration along the three cardinal axes are shown as a maximum displacement projection overlaid with 2D projections of the reconstructed axon skeletons. Rows indicate the viewing direction, columns indicate the displacement component color-coded in the heatmaps.

## REFERENCES

Agronskaia, A. V., J. A. Valentijn, L. F. van Driel, C. T. W. M. Schneijdenberg, B. M. Humbel, P. M. P. van Bergen en Henegouwen, A. J. Verkleij, A. J. Koster and H. C. Gerritsen (2008). “Integrated fluorescence and transmission electron microscopy.” Journal of Structural Biology 164(2): 183–189.

Anderson, J. C., R. J. Douglas, K. A. C. Martin and J. C. Nelson (1994). “Map of the synapses formed with the dendrites of spiny stellate neurons of cat visual cortex.” Journal of Comparative Neurology 341 (1): 25–38.

Anderson, J. C., R. J. Douglas, K. A. C. Martin and J. C. Nelson (1994). “Synaptic output of physiologically identified spiny stellate neurons in cat visual cortex.” Journal of Comparative Neurology 341(1): 16–24.

Atasoy, D., J. N. Betley, W.-P. Li, H. H. Su, S. M. Sertel, L. K. Scheffer, J. H. Simpson, R. D. Fetter and S. M. Sternson (2014). “A genetically specified connectomics approach applied to long-range feeding regulatory circuits.” Nature Neuroscience 17(12): 1830–1839.

Berck, M. E., A. Khandelwal, L. Claus, L. Hernandez-Nunez, G. Si, C. J. Tabone, F. Li, J. W. Truman, R. D. Fetter, M. Louis, A. D. T. Samuel and A. Cardona (2016). “The wiring diagram of a glomerular olfactory system.” eLife 5.

Berning, M., Kevin M. Boergens and M. Helmstaedter (2015). “SegEM: Efficient Image Analysis for High-Resolution Connectomics.” Neuron 87(6): 1193–1206.

Bishop, D., I. Nikić, M. Brinkoetter, S. Knecht, S. Potz, M. Kerschensteiner and T. Misgeld (2011). “Near-infrared branding efficiently correlates light and electron microscopy.” Nature Methods 8(7): 568–570.

Bock, D. D., W.-C. A. Lee, A. M. Kerlin, M. L. Andermann, G. Hood, A. W. Wetzel, S. Yurgenson, E. R. Soucy, H. S. Kim and R. C. Reid (2011). “Network anatomy and in vivo physiology of visual cortical neurons.” Nature 471 (7337): 177–182.

Boergens, K. M., M. Berning, T. Bocklisch, D. Bräunlein, F. Drawitsch, J. Frohnhofen, T. Herold, P. Otto, N. Rzepka, T. Werkmeister, D. Werner, G. Wiese, H. Wissler and M. Helmstaedter (2017). “webKnossos: efficient online 3D data annotation for connectomics.” Nature Methods 14(7): 691–694.

Briggman, K. L., M. Helmstaedter and W. Denk (2011). “Wiring specificity in the direction-selectivity circuit of the retina.” Nature 471 (7337): 183–188.

Colonnier, M. (1964). “Experimental Degeneration in the Cerebral Cortex.” Journal of anatomy 98: 47–53.

Costa, N. M. d. and K. A. C. Martin (2011). “How Thalamus Connects to Spiny Stellate Cells in the Cat’s Visual Cortex.” The Journal of Neuroscience 31(8): 2925–2937.

D’Souza, R. D., A. M. Meier, P. Bista, Q. Wang and A. Burkhalter (2016). “Recruitment of inhibition and excitation across mouse visual cortex depends on the hierarchy of interconnecting areas.” eLife 5.

Denk, W. and H. Horstmann (2004). “Serial Block-Face Scanning Electron Microscopy to Reconstruct Three-Dimensional Tissue Nanostructure.” PLoS Biol 2(11): e329.

Faulk, P. W. and M. G. Taylor (1971). “An immunocolloid method for the electron microscope.” Immunochemistry 8(11): 1081–1083.

Fua, P. and G. W. Knott (2015). “Modeling brain circuitry over a wide range of scales.” Frontiers in Neuroanatomy 9.

Grabenbauer, M., W. J. C. Geerts, J. Fernadez-Rodriguez, A. Hoenger, A. J. Koster and T. Nilsson (2005). “Correlative microscopy and electron tomography of GFP through photooxidation.” Nature Methods 2(11): 857–862.

Gray, E. G. and L. H. Hamlyn (1962). “Electron microscopy of experimental degeneration in the avian optic tectum.” Journal of Anatomy 96(Pt 3): 309–316.305.

Hamos, J. E., S. C. Van Horn, D. Raczkowski, D. J. Uhlrich and S. M. Sherman (1985). “Synaptic connectivity of a local circuit neurone in lateral geniculate nucleus of the cat.” Nature 317(6038): 618–621.

Hayworth, K. J., C. S. Xu, Z. Lu, G. W. Knott, R. D. Fetter, J. C. Tapia, J. W. Lichtman and H. F. Hess (2015). “Ultrastructurally smooth thick partitioning and volume stitching for large-scale connectomics.” Nature Methods 12(4): 319–322.

Helmstaedter, M. (2013). “Cellular-resolution connectomics: challenges of dense neural circuit reconstruction.” Nature Methods 10(6): 501–507.

Helmstaedter, M., K. L. Briggman and W. Denk (2011). “High-accuracy neurite reconstruction for high-throughput neuroanatomy.” Nature Neuroscience 14(8): 1081–1088.

Helmstaedter, M., K. L. Briggman, S. C. Turaga, V. Jain, H. S. Seung and W. Denk (2013). “Connectomic reconstruction of the inner plexiform layer in the mouse retina.” Nature 500(7461): 168–174.

Heymann, J. A. W., M. Hayles, I. Gestmann, L. A. Giannuzzi, B. Lich and S. Subramaniam (2006). “Site-specific 3D imaging of cells and tissues with a dual beam microscope.” Journal of structural biology 155(1): 63–73.

Horikawa, K. and W. E. Armstrong (1988). “A versatile means of intracellular labeling: injection of biocytin and its detection with avidin conjugates.” Journal of Neuroscience Methods 25(1): 1–11.

Horn, B. K. P. (1987). “Closed-form solution of absolute orientation using unit quaternions.” JOSA A 4(4): 629–642.

Hua, Y., P. Laserstein and M. Helmstaedter (2015). “Large-volume en-bloc staining for electron microscopy-based connectomics.” Nature Communications 6: 7923.

Joesch, M., D. Mankus, M. Yamagata, A. Shahbazi, R. Schalek, A. Suissa-Peleg, M. Meister, J. W. Lichtman, W. J. Scheirer and J. R. Sanes (2016). “Reconstruction of genetically identified neurons imaged by serial-section electron microscopy.” eLife 5: e15015.

Kasthuri, N., K. J. Hayworth, D. R. Berger, R. L. Schalek, J. A. Conchello, S. Knowles-Barley, D. Lee, A. Vázquez-Reina, V. Kaynig, T. R. Jones, M. Roberts, J. L. Morgan, J. C. Tapia, H. S. Seung, W. G. Roncal, J. T. Vogelstein, R. Burns, D. L. Sussman, C. E. Priebe, H. Pfister and J. W. Lichtman (2015). “Saturated Reconstruction of a Volume of Neocortex.” Cell 162(3): 648–661.

Knott, G., H. Marchman, D. Wall and B. Lich (2008). “Serial Section Scanning Electron Microscopy of Adult Brain Tissue Using Focused Ion Beam Milling.” The Journal of Neuroscience 28(12): 2959–2964.

Knott, G. W., A. Holtmaat, J. T. Trachtenberg, K. Svoboda and E. Welker (2009). “A protocol for preparing GFP-labeled neurons previously imaged in vivo and in slice preparations for light and electron microscopic analysis.” Nature Protocols 4(8): 1145–1156.

Lam, S. S., J. D. Martell, K. J. Kamer, T. J. Deerinck, M. H. Ellisman, V. K. Mootha and A. Y. Ting (2015). “Directed evolution of APEX2 for electron microscopy and proximity labeling.” Nature Methods 12(1): 51–54.

Lee, S., G. Wolberg and S.-Y. Shin (1997). “Scattered data interpolation with multilevel B-splines.” IEEE Transactions on Visualization and Computer Graphics 3(3): 228–244.

Lee, W.-C. A., V. Bonin, M. Reed, B. J. Graham, G. Hood, K. Glattfelder and R. C. Reid (2016). “Anatomy and function of an excitatory network in the visual cortex.” Nature 532(7599): 370–374.

Lin, T.-Y., J. Luo, K. Shinomiya, C.-Y. Ting, Z. Lu, I. A. Meinertzhagen and C.-H. Lee (2016). “Mapping Chromatic Pathways in the Drosophila Visual System.” The Journal of comparative neurology 524(2): 213–227.

Liv, N., A. C. Zonnevylle, A. C. Narvaez, A. P. J. Effting, P. W. Voorneveld, M. S. Lucas, J. C. Hardwick, R. A. Wepf, P. Kruit and J. P. Hoogenboom (2013). “Simultaneous Correlative Scanning Electron and High-NA Fluorescence Microscopy.” PLOS ONE 8(2): e55707.

Livet, J., T. A. Weissman, H. Kang, R. W. Draft, J. Lu, R. A. Bennis, J. R. Sanes and J. W. Lichtman (2007). “Transgenic strategies for combinatorial expression of fluorescent proteins in the nervous system.” Nature 450(7166): 56–62.

Maco, B., A. Holtmaat, M. Cantoni, A. Kreshuk, C. N. Straehle, F. A. Hamprecht and G.W. Knott (2013). “Correlative In Vivo 2 Photon and Focused Ion Beam Scanning Electron Microscopy of Cortical Neurons.” PLOS ONE 8(2): e57405.

Mao, T., D. Kusefoglu, Bryan M. Hooks, D. Huber, L. Petreanu and K. Svoboda (2011). “Long-Range Neuronal Circuits Underlying the Interaction between Sensory and Motor Cortex.” Neuron 72(1): 111–123.

Maranto, A. R. (1982). “Neuronal mapping: a photooxidation reaction makes Lucifer yellow useful for electron microscopy.” Science (New York, N.Y.) 217(4563): 953–955.

Markram, H., J. Lübke, M. Frotscher, A. Roth and B. Sakmann (1997). “Physiology and anatomy of synaptic connections between thick tufted pyramidal neurones in the developing rat neocortex.” The Journal of Physiology 500(2): 409–440.

Martell, J. D., T. J. Deerinck, Y. Sancak, T. L. Poulos, V. K. Mootha, G. E. Sosinsky, M. H. Ellisman and A. Y. Ting (2012). “Engineered ascorbate peroxidase as a genetically encoded reporter for electron microscopy.” Nature Biotechnology 30(11): 1143–1148.

Micheva, K. D. and S. J. Smith (2007). “Array tomography.” Neuron 55(1): 25–36.

Murphy, G. E., K. Narayan, B. C. Lowekamp, L. M. Hartnell, J. A. W. Heymann, J. Fu and S. Subramaniam (2011). “Correlative 3D imaging of Whole Mammalian Cells with Light and Electron Microscopy.” Journal of structural biology 176(3): 268–278.

Pallotto, M., P. V. Watkins, B. Fubara, J. H. Singer and K. L. Briggman (2015). “Extracellular space preservation aids the connectomic analysis of neural circuits.” eLife 4.

Rah, J.-C., E. Bas, J. Colonell, Y. Mishchenko, B. Karsh, R. D. Fetter, E. W. Myers, D. B. Chklovskii, K. Svoboda, T. D. Harris and J. T. R. Isaac (2013). “Thalamocortical input onto layer 5 pyramidal neurons measured using quantitative large-scale array tomography.” Frontiers in Neural Circuits 7.

Rueckert, D., L. I. Sonoda, C. Hayes, D. L. G. Hill, M. O. Leach and D. J. Hawkes (1999). “Nonrigid registration using free-form deformations: application to breast MR images.” IEEE Transactions on Medical Imaging 18(8): 712–721.

Saalfeld, S., A. Cardona, V. Hartenstein and P. Tomančák (2009). “CATMAID: collaborative annotation toolkit for massive amounts of image data.” Bioinformatics 25(15): 1984–1986.

Schalek, R., N. Kasthuri, K. Hayworth, D. Berger, J. Tapia, J. Morgan, S. Turaga, E. Fagerholm, H. Seung and J. Lichtman (2011). “Development of High-Throughput, High-Resolution 3D Reconstruction of Large-Volume Biological Tissue Using Automated Tape Collection Ultramicrotomy and Scanning Electron Microscopy.” Microscopy and Microanalysis; Cambridge 17(S2): 966–967.

Serradell, E., P. Glowacki, J. Kybic, F. Moreno-Noguer and P. Fua (2012). Robust non-rigid registration of 2D and 3D graphs. 2012 IEEE Conference on Computer Vision and Pattern Recognition.

Serradell, E., M. A. Pinheiro, R. Sznitman, J. Kybic, F. Moreno-Noguer and P. Fua (2015). “Non-Rigid Graph Registration using Active Testing Search.” 14.

Shahidi, R., E. A. Williams, M. Conzelmann, A. Asadulina, C. Verasztó, S. Jasek, L. A. Bezares-Calderón and G. Jékely (2015). “A serial multiplex immunogold labeling method for identifying peptidergic neurons in connectomes.” eLife 4.

Shu, X., V. Lev-Ram, T. J. Deerinck, Y. Qi, E. B. Ramko, M. W. Davidson, Y. Jin, M.H. Ellisman and R. Y. Tsien (2011). “A Genetically Encoded Tag for Correlated Light and Electron Microscopy of Intact Cells, Tissues, and Organisms.” PLoS Biol 9(4): e1001041.

Staffler, B., M. Berning, K. M. Boergens, A. Gour, P. v. d. Smagt and M. Helmstaedter (2017). “SynEM, automated synapse detection for connectomics.” eLife 6: e26414.

Stepanyants, A., L. M. Martinez, A. S. Ferecskó and Z. F. Kisvárday (2009). “The fractions of short- and long-range connections in the visual cortex.” Proceedings of the National Academy of Sciences of the United States of America 106(9): 3555–3560.

Tsang, T. K., E. A. Bushong, D. Boassa, J. Hu, B. Romoli, S. Phan, D. Dulcis, C.-Y. Su and M. H. Ellisman (2018). “High-quality ultrastructural preservation using cryofixation for 3D electron microscopy of genetically labeled tissues.” eLife 7: e35524.

Wanner, A. A., C. Genoud, T. Masudi, L. Siksou and R. W. Friedrich (2016). “Dense EM-based reconstruction of the interglomerular projectome in the zebrafish olfactory bulb.” Nature Neuroscience.

Xu, C. S., K. J. Hayworth, Z. Lu, P. Grob, A. M. Hassan, J. G. García-Cerdán, K. K. Niyogi, E. Nogales, R. J. Weinberg and H. F. Hess (2017). “Enhanced FIB-SEM systems for large-volume 3D imaging.” eLife 6.

